# Next Generation Neuropeptide Y Receptor Small Molecule Agonists Inhibit Mosquito Biting Behavior

**DOI:** 10.1101/2024.02.28.582529

**Authors:** Emely V. Zeledon, Leigh A. Baxt, Tanweer A. Khan, Mayako Michino, Michael Miller, David J. Huggins, Caroline S. Jiang, Leslie B. Vosshall, Laura B. Duvall

## Abstract

Female *Aedes aegypti* mosquitoes can spread disease-causing pathogens when they bite humans to obtain blood nutrients required for egg production. Following a complete blood meal, host-seeking is suppressed until eggs are laid. Neuropeptide Y-like Receptor 7 (NPYLR7) plays a role in endogenous host-seeking suppression and previous work identified small molecule NPYLR7 agonists that suppress host-seeking and blood feeding when fed to mosquitoes at high micromolar doses. Using structure activity relationship analysis and structure-guided design we synthesized 128 compounds with similarity to known NPYLR7 agonists. Although *in vitro* potency (EC_50_) was not strictly predictive of *in vivo* effect, we identified 3 compounds that suppressed blood feeding from a live host when fed to mosquitoes at a 1 μM dose, a 100-fold improvement over the original reference compound. Exogenous activation of NPYLR7 represents an innovative vector control strategy to block mosquito biting behavior and prevent mosquito/human host interactions that lead to pathogen transmission.

## Introduction

Female *Aedes aegypti* mosquitoes are innately attracted to find and bite human hosts to obtain blood protein required for egg development. However, host-seeking behavior is regulated by the female’s internal state and is naturally suppressed after a full meal of blood, during egg development (Judson CL, 1967). Blood nutrients are required for sustained suppression; although female mosquitoes will engorge on non-nutritive saline meals that cannot support egg development these females return to high levels of host-seeking when abdominal distensions wears off roughly 24 hours later (Duvall et al., 2019; Galun, 1963; Klowden and Lea, 1979b). Previous work implicates abdominal mechanosensors in mediating short-term suppression and neuropeptide pathways in mediating sustained, days-long, suppression (Brown et al., 1994; Christ et al., 2018; Davis, 1984; Duvall et al., 2019; Klowden, 1981; Klowden and Lea, 1979a, 1979b; Liesch et al., 2013). Neuropeptide Y-related pathways regulate hunger and satiety in many organisms (Colmers and Wahlestedt, 1993; De Bono and Bargmann, 1998; Inui, 1999; Kuenzel et al., 1987; Wu et al., 2003) and we recently identified *Aedes aegypti* Neuropeptide Y-like Receptor 7 (NPYLR7) as a key regulator of host-seeking after a blood meal (Duvall et al., 2019). After blood feeding, NPYLR7 activation acts as a satiety signal and suppresses attraction to hosts. Pharmacological activation of NPYLR7 inhibits biting and blood feeding even in the absence of blood nutrients. Conversely, female mosquitoes with genetically or pharmacologically disrupted NPYLR7 signaling continue to host-seek inappropriately after a blood meal. Exogenous activation of feeding-related neuropeptide receptors in mosquitoes represents a novel approach to block their attraction to humans by exploiting the pathways that naturally suppress the drive to bite.

Here, we used structure activity relationship analysis and structure-guided design to identify novel small molecule NPYLR7 agonists with improved *in vitro* and *in vivo* potency compared to compounds identified in a previous high-throughput small molecule screen of 265,211 compounds (Duvall et al., 2019). We synthesized 128 new compounds and characterized their *in vitro* potency using an HEK cell-based assay to evaluate NPYLR7 activation. To identify those with *in vivo* activity, we tested 30 compounds in a host-seeking screening assay and subsequently identified 3 compounds that significantly reduced blood feeding from a live host when delivered to mosquitoes at a 1 μM, a dose 100-fold lower than that used in our original report (Duvall et al., 2019).

Although 3 out of 30 compounds tested *in vivo* suppressed host-seeking behavior, *in vitro* potency was not highly predictive of *in vivo* efficacy. This disconnect highlights the need for intermediate assays to span the gap between cell-based and behavioral assays in mosquitoes. This work is important because it identifies new highly potent compounds that block mosquito blood feeding by targeting a neuropeptide receptor that regulates mosquito attraction to humans through a conserved satiety pathway.

## Results

We previously showed that pharmacological activation of NPYLR7 suppressed *Aedes aegypti* mosquito host-seeking when delivered in a non-nutritive saline meal and we identified small molecule NPYLR7 agonists that induce host-seeking suppression independent of nutrient consumption (Figure 1A). However, the most potent of these compounds (TDI-012631) had a half-maximal effective concentration (EC_50_) in activating NPYLR7 *in vitro* of 19.6 μM and was behaviorally active only when fed at high micromolar doses (> 30 μM). We therefore set out to design and evaluate analogs of this compound to identify those with improved potency. We generated a docking model of TDI-012631 bound to NPYLR7 to identify protein-ligand interactions that could inform analog design. The docking model showed that the quinazoline core occupies a hydrophobic pocket in the orthosteric binding site formed by transmembrane helices 3, 5, and 6. The guanidine substituent is stabilized by a salt bridge interaction with the Glu198 side chain in EL2 loop, while the 4-position methyl group is pointed toward Gln122^3.32^ and 7-position methoxy group is oriented toward Phe218^5.47^ (Figure 1B). This docking model is in agreement with early structure-activity relationship data for close analogs of TDI-012631, where the quinazoline core and guanidine appear to form a minimum pharmacophore. Indeed, replacement of the guanidine group with an amine (compound TDI-012610) causes a loss of potency. Newly synthesized analogs were then tested using a calcium-based HEK293T assay to determine their *in vitro* potency compared to FMRFa3, an endogenous peptide ligand of NPYLR7 (Figure 1 C and D). We used the original reference compound, TDI-012631, as a benchmark for potency. Based on the *in vitro* assay, compounds were grouped by EC_50_ value into those that showed low sensitivity or potency (> 100 μM), those with *in vitro* EC_50_ values < 100 μM but > 4.11 μM (TDI-012631, reference compound) and those with EC_50_ values < 4.11 μM.

**Figure 1:**
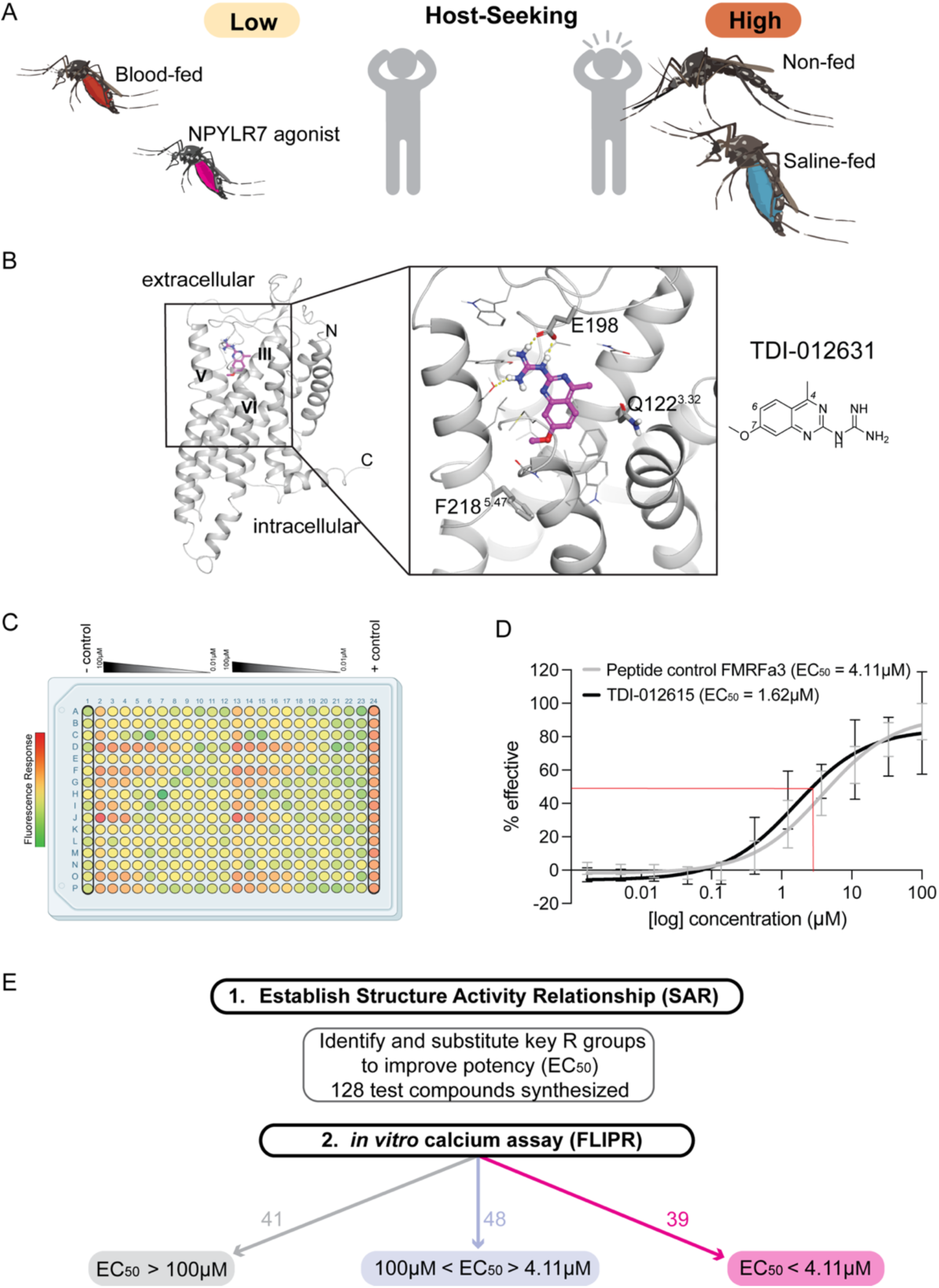
Structure-guided design to synthesize novel NPYLR7 agonists. (A) Representation of *Aedes aegypti* host-seeking behavior by feeding status. (B) NPYLR7 predicted structure model with compound TDI-012631 bound (left) and predicted side chain interactions (right). (C) Representative 384 well plate layout from *in vitro* screen displaying Raw Fluorescent Units (RFU) corresponding to test compounds ranging from concentration of 100 - 0 μM from left to right. Negative control (column 1) is measured as response to assay buffer alone and positive control (column 24) is response to 10 μM dose of FMRFa3, an endogenous peptide activator of NPYLR7. (D) Semilogarithmic curves of compound TDI-012615 (EC_50_ = 1.62 μM, black line) and FMRFa3 (peptide control) sigmoidal curve (EC_50_ = 4.11 μM, gray line). (E) Outline of *in vitro* screening for 128 newly synthesized NPYLR7 agonists binned according to *in vitro* EC_50_.

Analogs of TDI-012631 were designed by maintaining the quinazoline core and the guanidine substituent that formed critical interactions in the docking model and modifying the substituents around the core at the 4 (R2), 6 (R1), and 7 (R) positions (Figure 2A). Bulkier groups were explored for R to extend into the deeper pocket, while hydrogen-bond donating NHCH_3_ group was introduced for R2 to interact with Gln122^3.32^ side chain (Figure 2A). Over a hundred compounds were designed and synthesized. To identify compounds with *in vivo* activity, thirty compounds were selected for testing in host-seeking assays. This group included compounds with predicted EC_50_ values ranging from 39.3 μM to 1.92 nM as well as a negative control compound from the > 100 μM group (Figure 2B).

**Figure 2:**
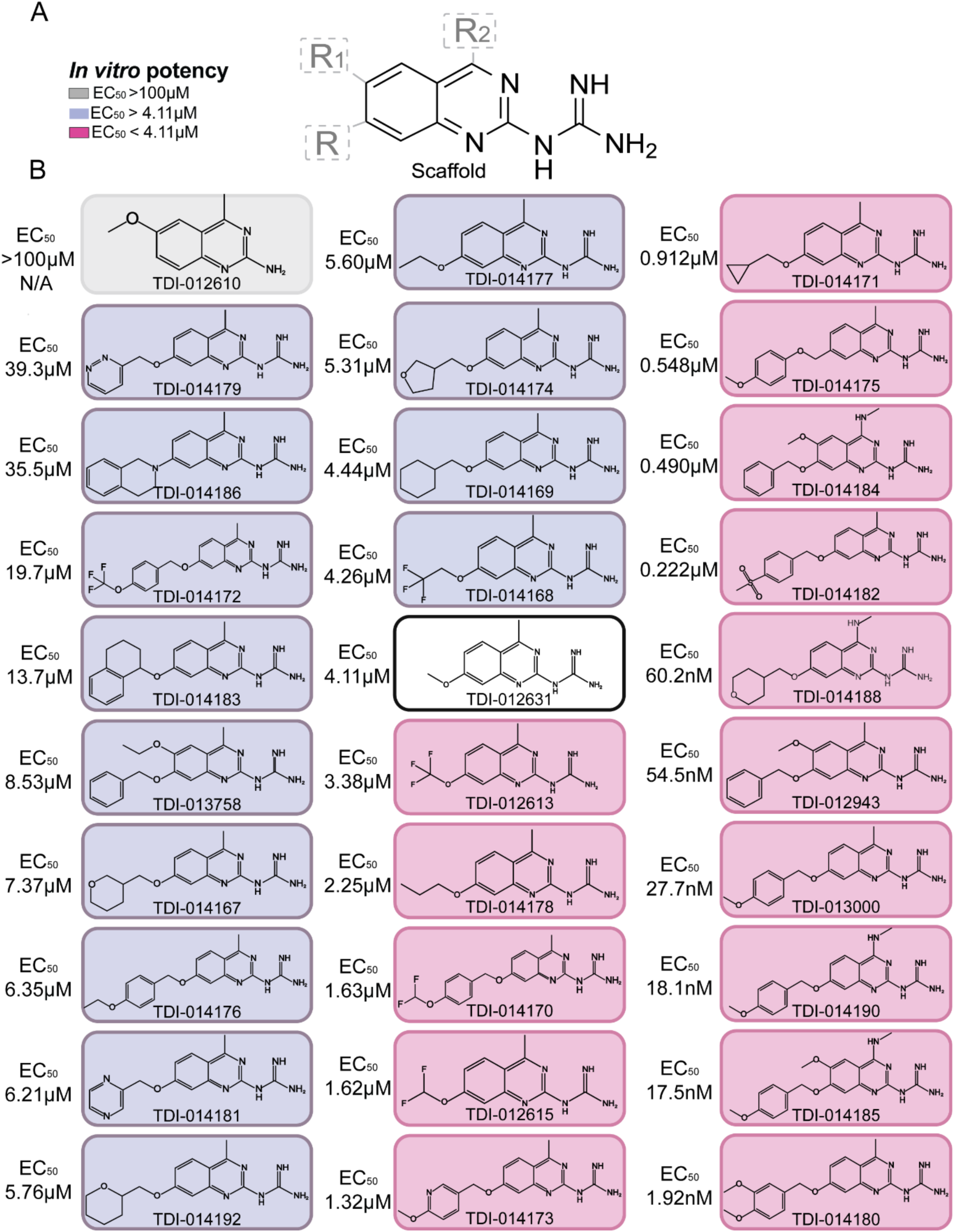
Small molecule NPYLR7 agonists tested *in vivo*. (A) Lead compound scaffold with conserved quinazoline core and R groups tested for substitutions. Shading in legend indicates EC_50_ in (B). (B) Chemical structures with corresponding *in vitro* EC_50_ of each compound tested in behavioral assays arranged with the highest value at the top left to the lowest value at bottom right. TDI-012631 (center, black border) is the initial reference compound.

To prioritize candidates with highest levels of *in vivo* efficacy we performed a screening “Miniport olfactometer” assay that allowed us to test mosquito host-seeking behavior in 4 conditions in parallel (Figure 3A). In this assay we provided two host cues, CO_2_ and human odor collected on a worn nylon stocking. Animals were scored as attracted if they flew from the starting canister into the attraction trap next to the source of the host cues. Animals were fed each compound at a 1 μM dose in non-nutritive saline 2 days prior to testing, then scored and weighed to ensure that compounds did not affect meal palatability or consumption (Data S1).

**Figure 3:**
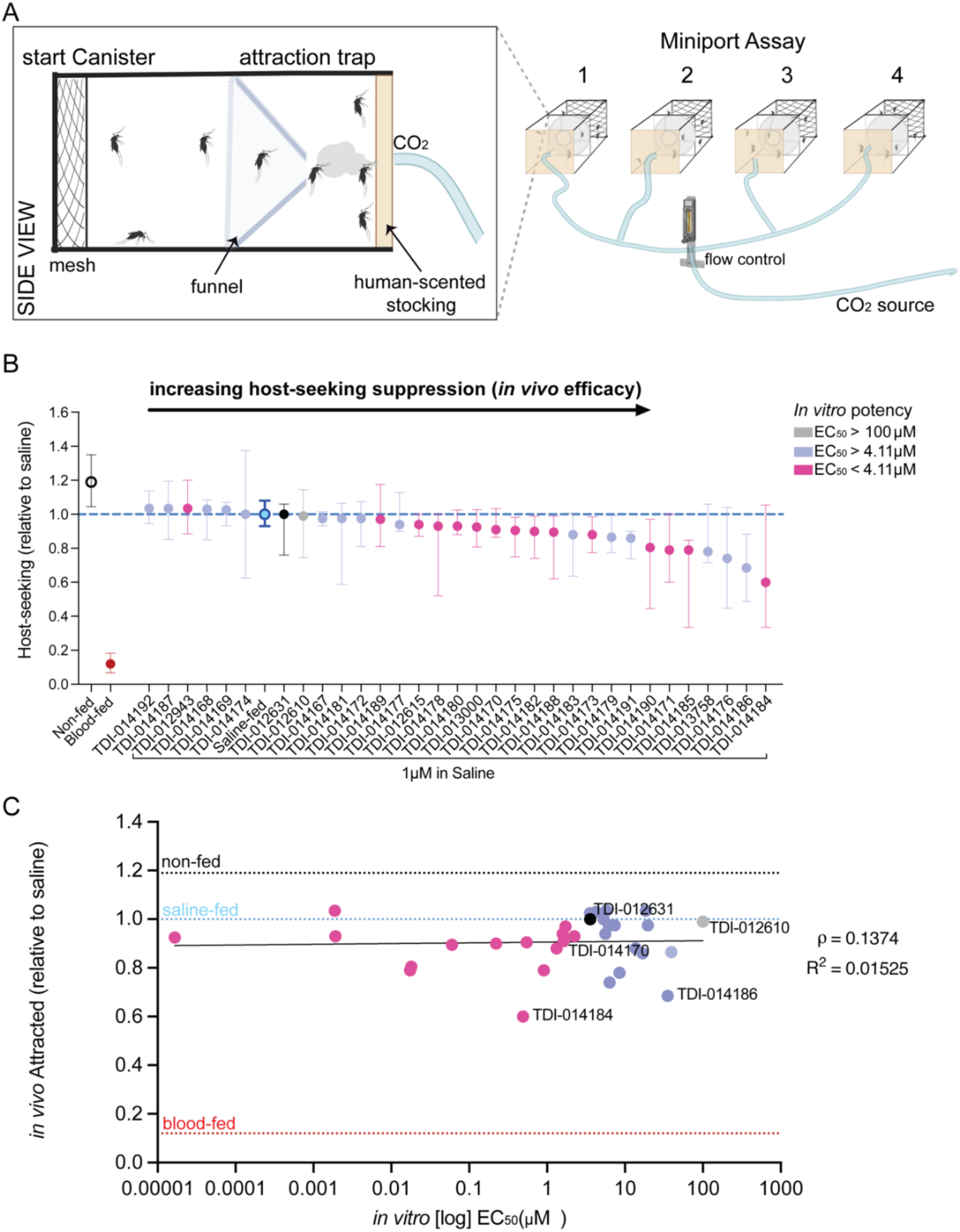
Miniport olfactometer screening assay to identify compounds that suppress attraction to human host cues. (A) Schematic of Miniport Olfactometer host-seeking assay. Inset depicts start canister and attraction trap modular components for each experiment. Mosquitoes not drawn to scale. (B) Host-seeking suppression relative to saline (test compound % host seeking/ average matched saline % host-seeking). (Median with interquartile range, n = 4 – 65 replicates, 15-20 females/replicate). (C) Correlation of *in vitro* potency to *in vivo* host seeking suppression in *Aedes aegypti*. Spearman correlation coefficient (ρ) = 0.1374, p > 0.05, slope = 0.002942, R^2^ = 0.01525.

Animals were allowed to recover for 2 days before host-seeking assays were performed to ensure that suppression was not attributable to abdominal distension from meal consumption. Using the Miniport olfactometer assay we identified compounds that suppressed host-seeking compared to saline alone (Figure 3B). Consistent with previous work, the original lead compound TDI-012631, was not active at a 1 μM dose. We plotted the relationship between *in vitro* EC_50_ and *in vivo* efficacy in the Miniport assay and found that, although we identified compounds that were more effective than TDI-012631 there was no predictive relationship between *in vitro* EC_50_ and *in vivo* efficacy (Figure 3C).

We next asked whether compounds that suppressed attraction to host cues in the Miniport olfactometer assay could also block blood feeding from a live host using a “Mouse in Cage” assay in which mosquitoes are fed test compounds and presented with an opportunity to blood feed from an anesthetized mouse 2 days later. Non blood-fed females were robustly attracted to the mouse and blood fed at high rates (91.0 ± 1.92%), while females that were naturally suppressed after a meal of sheep blood rarely fed (7.0 ± 1.21%) (Data S1). At the end of each experiment mosquitoes were collected and scored for the presence of fresh blood in their abdomen, indicating that they successfully fed on the blood of the mouse (Figure 4A). We replicated the finding that TDI-012631 suppressed blood feeding when fed to mosquitoes at a 100 μM dose (Figure 4B) (Duvall et al., 2019). Although TDI-014170 showed high *in vitro* potency (EC_50_ = 1.63 μM), this compound did not reduce host-seeking in our Miniport olfactometer screening assay nor did it significantly reduce biting in the Mouse in Cage assay when fed at a 1 μM dose (Figure 4C). However, we identified 3 novel small molecule NPYLR7 agonists that reduced biting behavior when fed at a 1 μM dose (Figure 4 D – F). These included TDI-014188 and TDI-014186, which were the 2 compounds that showed the largest effect in the Miniport host-seeking assay (Figure 3B). Although two of the three active compounds TDI-014184 (EC_50_ = 0.490 μM) (Figure 4F) and TDI-014188 (EC_50_ = 60.2 nM) (Figure 4D) showed significantly improved *in vitro* potency compared to the reference compound, surprisingly, (TDI-014186) (Figure 4E) effectively suppressed biting despite showing lower *in vitro* potency (EC_50_ = 35.5 μM) compared to the reference compound.

**Figure 4:**
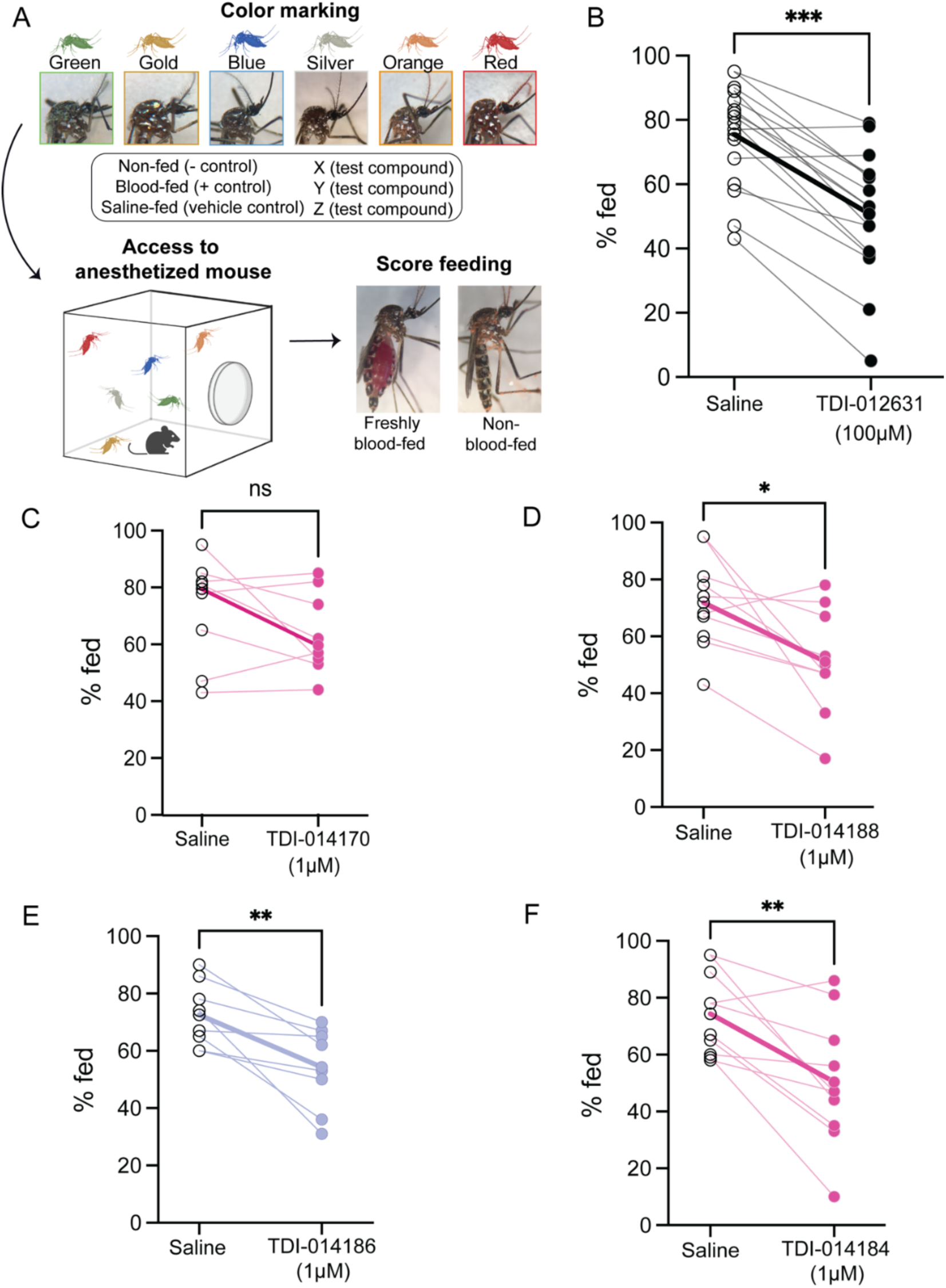
Novel NPYLR7 agonists suppress blood feeding from a live host when fed at 1 μM dose. (A)Schematic of “mouse in cage” biting assay workflow depicting powder color assignment, exposure to mouse, and scoring of fresh blood in the abdomen. (B – F) Percentage of females that freshly blood fed on an anesthetized mouse after 15 minute exposure. Females were fed the indicated meal 48 hours prior to the experiment. (C,D,F) are high potency and (E) is a medium potency *in vitro* compound (See Figure 2B). Bold lines represent saline group mean compared to test group mean. 15 – 20 females per replicate. Mann-Whitney test: (B) n = 13 replicates, ***p = 0.0003, (C) n= 9 replicates, ns, not significant, p = 0.2868, (D) n= 11 replicates, *p = 0.0120). (E) n= 9 replicates, ** p = 0.0097, (F) n= 11 replicates; ** p = 0.0055.

## Discussion

Multiple mosquito genera contribute to the spread of human disease. *Aedes* mosquitoes can transmit the dangerous arboviruses yellow fever, dengue, Zika, and chikungunya (Bhatt et al., 2013; Weaver et al., 2013). The emergence and geographical spread of these viruses are critical concerns for global public health (Bhatt et al., 2013; Kraemer et al., 2015; Ryan et al., 2019). Although much research has been dedicated to developing vaccines and prophylactic or therapeutic drugs to treat and prevent these diseases there are currently no drugs to treat the viruses spread by *Aedes aegypti*. Although there is an effective vaccine against the yellow fever virus and there have been significant improvements in recent dengue vaccination strategies (Adams et al., 2021), preventing mosquito bites remains essential for reducing disease transmission. Current strategies to control mosquito populations rely on toxic pesticides that decline in efficacy as mosquito populations rapidly develop resistance (Moyes et al., 2021). More recently-developed approaches involve the release of mosquitoes rendered either sterile or unable to transmit pathogens (Blumberg et al., 2016; Lees et al., 2015) or releasing transgenic animals with altered immune or reproductive function (Hammond et al., 2016; Macias et al., 2017; Parihar et al., 2020). However, ethical, environmental, and regulatory concerns remain issues in deployment of such transgenic mosquitoes. Although each of these approaches has shown some success, there remains major unmet need to develop innovative and complementary strategies for integrated mosquito control.

The most prevalent chemical control methods include synthetic insecticides or repellents and previous studies have primarily focused on determining pharmacokinetic properties of these compounds in non-target vertebrates at doses lethal to the target insect (González-Morales et al., 2023; Qiu et al., 1997). Lethality contributes to evolution of resistance by selecting for individuals carrying resistance alleles that can escape lethality (Liu, 2015; Su et al., 2019). Most insecticides are limited to a few chemical classes with similar mechanisms of action and the World Health Organization has urged the development of new mosquito control techniques that exploit novel chemical classes (World Health Organization, 2012). Increasingly, new strategies have focused on non-lethal methods of pest control (Savino et al., 2018; Witmer et al., 2017). This category includes the small molecule NPYLR7 agonists tested in this study, which reduce the drive to bite without killing the mosquito. However, there is a need for relevant drug discovery assays in insects to characterize the utility and potential for development of non-lethal chemicals. In traditional drug discovery and development, *in vitro* assays to determine potency are normally followed by assays to determine target engagement in cells and pharmacokinetic/pharmacodynamic (PK/PD) properties to evaluate how compounds are absorbed, metabolized and eliminated from specific tissues in the body. There are examples of insect models for drug discovery; silkworms (*Bombyx mori*) have been used as a model for drug toxicity and show responses to hepatotoxic drugs that are consistent with mammalian models (Hamamoto et al., 2009; He et al., 2020). More recent work has established novel models for pharmacokinetic assays in mosquitoes by delivering ivermectin and cytochrome P450 modulators in blood, using liquid chromatography with tandem mass spectrometry (LC-MS/MS) to quantify clearance rates after feeding, and modeling primary pharmacokinetic parameters and drug/drug interactions. Ivermectin clearance kinetics differ between mosquitoes compared to mammals, and individual P450 modulators were eliminated with differing kinetics in mosquitoes (Duthaler et al., 2021). However, detection remains a limiting factor. Drug concentrations are higher than those that can be obtained in the blood of humans receiving a regular dose of ivermectin and because whole mosquito specimens were required for analysis it is not yet possible to achieve tissue-specific resolution of drug occupancy/clearance. We found that *in vitro* EC_50_ was not predictive of behavioral effect among the compounds tested in our assays and future work to characterize the clearance rates of the compounds described here may clarify the relationship between EC_50_ in our cell-based assay and behavioral effect. Compounds like TDI-014170 with high *in vitro* potency may be inactive *in vivo* because they lack bioavailability in the mosquito, or have a half-life that is too short to be captured when assays are performed 2 days after feeding. Compounds like TDI-014186 that suppressed host-seeking behavior despite modest *in vitro* EC_50_ may achieve *in vivo* potency through sequestration in specific target tissues.

NPY-like receptors are present in many insects including other blood-feeding arthropods (Garczynski et al., 2005; Gulia-Nuss et al., 2016). This suggests that NPY pathways may represent a conserved biological mechanism that could be targeted for the development of a more generalized strategy to suppress attraction to humans across multiple species of mosquitoes and ticks. To ensure that beneficial insects are not impacted, *in vitro* assays to identify compounds that show minimal cross-activation of related receptors in other insect species will be crucial. Delivering these compounds in traps baited with human odor will also ensure that beneficial insects are not targeted, a method is already in use in attractive toxic sugar baits (Fiorenzano et al., 2017; Krockel et al., 2006; Mukabana et al., 2012; Okumu et al., 2010).

By combining advanced molecular modelling, medicinal chemistry and precise deployment strategies, there is potential to develop targeted and environmentally responsible solutions for managing mosquito populations. Continued research into the structural aspects of these compounds and their receptor interactions will pave the way for the development of next-generation insect control agents with broader applicability and improved efficacy. Further study of the structural relationship between these compounds and *Aedes aegypti* NPYLR7 may help to identify new highly potent compounds and the rules by which they activate their cognate receptors.

## Supporting information

Supplemental methods for synthesis reactions

## Acknowledgements

We thank Gloria Gordon, Libby Mejia, and Ellen de Obaldia for assistance with mosquito rearing and Jim Petrillo for support with Miniport construction. Support for this project was provided by an Advanced Grant from the Robertson Therapeutic Development Fund, generously provided by the Robertson Foundation. The authors gratefully acknowledge the support to the project generously provided by the Sanders Tri-Institutional Therapeutics Discovery Institute (TDI), a 501(c)(3) organization. TDI receives financial support from Takeda Pharmaceutical Company, TDI’s parent institutes (Memorial Sloan Kettering Cancer Center, The Rockefeller University and Weill Cornell Medicine) and from a generous contribution from Mr. Lewis Sanders and other philanthropic sources. This work was partially supported by a Clinical and Translational Science Award (UL1 TR001866) from the National Center for Advancing Translational Sciences at the National Institutes of Health to C.S.J., a grant from NIGMS (R35 GM137888), a Beckman Young Investigator Award, a Pew Scholar in Biomedical Sciences Award, and a Klingenstein-Simons Fellowship Award in Neuroscience to L.B.D. L.B.V. is supported by the Howard Hughes Medical Institute.

## Author Contributions

Mayako Michino and D.H. carried out molecular modeling, T.K. and Mike Miller coordinated compound synthesis, and L.A.B. coordinated compound profiling and *in vitro* data analysis. C.S.J. provided statistical expertise for *in vivo* data analysis. E.V.Z. carried out all *in vivo* experiments. E.V.Z., L.B.D., and L.B.V. conceived the study, designed the figures, and wrote the paper, with input from all authors.

## Materials and Methods

### Molecular modelling for the NPYLR7 agonist series

The docking model for the NPYLR7 agonist series was generated based on a homology model of *Aedes aegypti* NPYLR7 and validated with rigorous FEP+ binding energy calculations (Abel et al., 2017). The homology model of *Aedes aegypti* NPYLR7 was obtained from the GPCR-I-Tasser homology modeling server (https://zhanggroup.org/GPCR-I-TASSER/) (Zhang et al., 2015). To predict the binding mode of reference compound TDI-012631, the compound was docked to the orthosteric binding site of NPYLR7 homology model using Glide SP, then the receptor-ligand complex was refined in a POPC lipid bilayer environment using Desmond molecular dynamics (MD) simulation for 120 ns at constant temperature of 300K in NPγT ensemble. The protein structure was prepared using the Protein Preparation Wizard in Maestro with default settings. The ligand structure was prepared using LigPrep in Maestro.

The protonated form of the ligand was selected, according to the predicted pKa of the guanidine moiety (pKa = 10.21 in Jaguar). The MD simulations were carried out using the OPLS3e force field (Harder et al., 2016). The final snapshot at 120 ns was minimized and then subjected to an absolute FEP+ calculation, (Khalak et al., 2021) which showed favorable binding energy (ΔG = -18.02 ± 0.24 kcal/mol). To further validate the docking model, relative FEP+ calculations were performed on a validation set of 13 compounds in the series having EC_50_ values ranging across 3 log units. The validation showed good agreement between the experimental and FEP+ predicted potencies (MUE = 1.29 kcal/mol; R^2^ = 0.57). The perturbation maps were automatically generated using the Mapper tool. Force Field Builder was employed to generate custom torsional parameters for ligand torsions that were not include in the default force field. FEP+ calculations were run for the default 5 ns. The Schrödinger Suite was used for protein and ligand preparations, docking, MD, and FEP+ calculations (release 2020-4, Maestro, Schrödinger LLC, New York, NY).

### Synthetic methods for analog preparation

Unless otherwise noted, the following pertain to the synthetic methods: all reactions are magnetically stirred; typical solvents (ethyl acetate, hexanes, dichloromethane, and methanol) are Fisher Optima grade; “concentrated to dryness” or “removal of the solvent” means evaporating the solvent from a solution or mixture using a rotary evaporator; flash chromatography is carried out on an Isco, Analogix, or Biotage automated chromatography system using a commercially available cartridge as the column. Columns are usually filled with silica gel as the stationary phase; preparative HPLC (or prep-HPLC) is carried out with commercial columns in a reverse phase manner (the stationary phase is hydrophobic). Typical solvent mixtures include A (water) and B (organic i.e. acetonitrile, methanol, etc.). Additives can also be used in the solvent mixture such as HCl, NH_4_HCO_3_, and formic acid. Details for individual synthesis reactions are provided in Supplemental Methods.

### *In vitro* Assay

The *in vitro* screening assay was adapted from (Duvall et al., 2019), and carried out at HD Biosciences (HDB, Shanghai) as follows: HEK293T (ThermoFisher Scientific) cells were grown in DMEM (high glucose, with glutamine), 10% Fetal Bovine Serum, 1% Pen/Strep, seeded in a 75 cm^2^ flask, and incubated at 37°C and 5% CO_2_. Cells were transiently transfected with 1 μg of each plasmid expressing GCaMP6s (Addgene #277314.1040753) (Chen et al., 2013), mouse Gqα15 (Addgene #40753) (Offermanns and Simon, 1995) and *Aedes aegypti* NPYLR7 (Addgene #52392) (Duvall et al., 2019) using Lipofectamine 2000 (Invitrogen) in 4mL of Opti-Mem (Invitrogen). This mixture was added to a plate after 20 minutes and incubated for 6 - 8 hours. Cells were then trypsinized, resuspended in phenol-free media, and plated in a 384 well plate (Greiner Bio-one) at a density of 20,000 cells per 40 μL, and incubated overnight (37°C, 5% CO_2_). Cells were imaged directly in phenol-free mediate. Plates were loaded into a Molecular Devices Fluorescent Imaging Plate Reader (FLIPR) with an excitation wavelength of 470 – 495 nm and an emitted wavelength of 515 – 575 nm. Plates were imaged every second for 5 minutes. After 30 seconds of baseline fluorescence recording, 10 μL of test compound in Reading Buffer ([Hanks’s Balanced Salt Solution (GIBCO) + 20 mM HEPES (Sigma-Aldrich)] was added. Concentrations tested ranged from 0 - 100 μM. Data was collected in raw fluorescence units (RFU).

### EC_50_ Calculations

Each test plate included replicates of 10 μM FMRFa3, an *Aedes aegypti* neuropeptide agonist of NPYLR7 used to calculate Hundred Percent Effect (HPE). The Zero Percent Effect (ZPE) control was calculated as response to buffer alone. Z’ Factor was calculated for each plate by 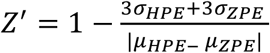. Plates with Z’ < 0.5 were excluded from analysis. Raw Fluorescent Units (RFUs) were converted to percent effect relative to the average of the ZPE and HPE. Percent Effect = (sample luminescence-average ZPE)/(average HPE-average ZPE)*100. Doses ranged from 0 to 100 μM, the highest 100 μM dose was excluded in replicates in which this dose showed aberrant responses beyond plateau (defined as 2 points with < 15% change).

Each replicate was plotted as percent effect vs log[concentration] and replicates with maximal responses between 20% and 250% were used for EC_50_ calculations. Relative EC_50_ was calculated using a [Agonist] versus response - variable slope four parameter model where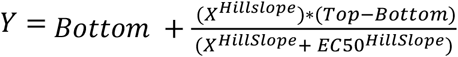. For compounds with multiple replicates, the relative EC_50_ of each replicate was averaged.

### Mosquito Rearing and Maintenance

*Aedes aegypti* (Orlando strain) were reared and maintained in a STERIS environmental room under the following conditions: 26 - 28°C, 80% humidity. Light cycles consisted of 14:10 hr (L:D). Eggs were hatched in hatching broth: 1 fish food tablet (TetraMin Tropical Tablets, Pet Mountain) crushed using a mortar and pestle brought to a volume of 850 mL diH2O and autoclaved for sterility). During larval stages (days 2 - 5), larvae were fed 2 tablets of fish food per day. Pupae were collected and allowed to eclose in 91 cm × 61 cm × 61 cm BugDorm cages (BioQuip Products). Adults were provided with cotton dental wicks (Richmond Dental) inserted into Boston clear round 60 mL glass bottle (ThermoFisher) filled with 10% sucrose (w/v). Adults were co-housed with siblings and allowed to mate freely for seven days post-eclosion. All behavioral experiments were performed using 14 - 21-day old females as in (Duvall et al., 2019). Males were removed prior to all behavioral experiments.

### Glytube feeding

Groups of 60 - 150 females were placed in 21.6 cm diameter x 16.5 cm height bucket cages (VWR) and fasted with access to water for 24 hours prior to feeding. Meals were delivered to females via Glytube membrane feeders as described (Costa-da-Silva et al., 2013). Meals consisted of 1.5 mL sheep blood (Hemostat Laboratories), saline (400 mM NaHCO_3_ + dih_2_O), or saline and test compound. Lyophilized compounds were stored at room temperatures until use. High concentration stocks (30 mM in 100% DMSO) (Sigma-Aldrich) were stored at -20°C and diluted in saline immediately prior to feeding. Once prepared, meals and glycerol heating elements were warmed in a 42°C water bath for 15 minutes to provide warmth to attract mosquitos to feed. ATP (1 mM final concentration) and test compounds (1 μM final concentration) were added prior to placing the meal on top of the mesh on the cage containing female mosquitos. Females were allowed 15 - 20 minutes to feed to repletion. Abdominal engorgement was scored by eye and confirmed by weighing females to ensure full feeding (Data S1). Females that did not feed to repletion were discarded. Females were returned to their original cage with a diH_2_O wick for 48 hours prior to behavioral testing. Lethality of each compound was measured by counting the number of dead females in each cage 24 hours post meal. Any compounds that resulted in > 50% death rate were scored as “high lethality” and excluded from behavioral testing.

### Behavior

#### Host Seeking Screening Assay (Miniport Olfactometer)

Miniport olfactometers were fabricated in-house as described (Duvall et al., 2019). Information regarding design, construction, and use can be found at https://github.com/VosshallLab/Miniport-Construction. Miniport canisters consisted of a 6’’ x 3’’ x 3’’ acrylic tube with a mesh screen to allow for air flow. Groups of 10 - 20 females were loaded into canisters 24 hours post meal. Females were left in canisters overnight to acclimate with two cotton balls soaked in diH_2_0 to prevent desiccation. Behavior trials began 48 hours post meal. Canisters were randomly assigned to attraction traps 1 - 4 and given 5 minutes to acclimate. The stimulus end of the trap was connected to a flowmeter that supplied 5% CO_2_ at a rate of 30 mL/min. Thirty seconds of CO_2_ flow was supplied to activate females prior to opening the sliding door. Once opened, the sliding door gave females access to the attraction trap baited with a human scented nylon stocking previously worn by the same experimenter for 8 -10 hours to collect body odor and stored in a plastic bag at -20°C until use. After 5 minutes, the sliding door was closed and attraction was scored as mosquitoes in attraction trap/total number mosquitoes. For each experiment data were normalized to the matched average saline attraction from the same experiment (% attracted/average saline attraction). Any dead mosquitoes were excluded from analysis. Non-fed females and blood-fed females served as environmental control groups while the saline-fed females served as the vehicle control each day. If non-fed or saline-fed females were less than 50% attracted or if blood-fed females were more than 20% attracted behavioral trials were halted and all data from that experimental day were discarded.

#### Biting Assay (Mouse in cage)

This assay was modified from (Duvall et al., 2019). Females were fed and scored as described above. 48 hours post meal, females were anesthetized in a 4°C cold room and aspirated into a petri dish (ThermoFisher Biosciences) with a randomly assigned color powder (Slice of the Moon; Chameleon Colors) to mark their treatment group. Females were then individually removed from the color powder, placed inside a 91 cm × 61 cm × 61 cm cage (BugDorm), and allowed 2 - 4 hours to recover. Each experiment consisted of a cage with 12 - 20 females in each treatment group including environmental (non-fed and blood-fed) and vehicle (saline-fed) control groups. The experiment began once an anesthetized mouse was introduced to the center of the cage and females were allowed 15 minutes to blood feed. The mouse was then removed, and the cage was placed at 4°C to anesthetize females for collection. Females were aspirated into a large glass petri dish (Pyrex) and scored under a dissection microscope for powder color and blood feeding status (Nikon SMZ1500). Females were scored as “blood-fed” if fresh blood was present in the abdomen. Percent biting was calculated as females freshly blood fed/total females in each respective treatment group.

#### Quantification and statistical analysis

All statistical analysis was performed using GraphPad Prism Version 10. Data collected as percentage of total are shown as median with range. Data collected as raw values are shown as mean ± SEM or mean ± SD. Details of statistical methods are reported in the figure legends.

### Data and software availability

Raw data are provided in Data S1. Cartoons in Figures 1, 3, and 4 were created with BioRender.com. Chemical structures in Figures 1 and 2 were generated with ChemDraw (version 21.0.0). The Schrödinger Suite was used for protein and ligand preparations, docking, MD, and FEP+ calculations (release 2020-4, Maestro, Schrödinger LLC, New York, NY).

### Human and animal ethics statement

Blood-feeding procedures and mosquito behavior with human scent collected on nylon were approved and monitored by The Rockefeller University Institutional Review Board (IRB protocol LVO-0652) and the Rockefeller University Institutional Animal Care and Use Committee (IACUC protocol 20068-H). Human subjects gave their written informed consent to participate in this study.

## Notes

### Competing Interest Statement

The authors have declared no competing interest.

### Summary of Updates

This version has been revised to update the NPYLR7 binding model shown in Figure 1B.

## References

Abel R, Wang L, Harder ED, Berne BJ, Friesner RA. 2017. Advancing Drug Discovery through Enhanced Free Energy Calculations. Acc Chem Res 50:1625–1632. doi:10.1021/acs.accounts.7b00083

Adams LE, Waterman S, Paz-Bailey G. 2021. Vaccination for Dengue Prevention. JAMA 327:817–818. doi:10.1001/jama.2021.23466

Bhatt S, Gething PW, Brady OJ, Messina JP, Farlow AW, Moyes CL, Drake JM, Brownstein JS, Hoen AG, Sankoh O, Myers MF, George DB, Jaenisch T, Wint GRW, Simmons CP, Scott TW, Farrar JJ, Hay SI, William Wint GR, Simmons CP, Scott TW, Farrar JJ, Hay SI, Wint GRW, Simmons CP, Scott TW, Farrar JJ, Hay SI. 2013. The Global Distribution and Burden of Dengue. Nature 496:504–507. doi:10.1038/nature12060

Blumberg BJ, Short SM, Dimopoulos G. 2016. Employing the Mosquito Microflora for Disease Control, Genetic Control of Malaria and Dengue. Elsevier Inc. doi:10.1016/B978-0-12-800246-9.00015-6

Brown MR, Klowden MJ, Crim JW, Young L, Shrouder LA, Lea AO. 1994. Endogenous Regulation of Mosquito Host-seeking Behavior by a Neuropeptide. J Insect Physiol 40:399–406. doi:10.1016/0022-1910(94)90158-9

Christ P, Hill SR, Schachtner J, Hauser F, Ignell R. 2018. Functional characterization of mosquito short neuropeptide F receptors. Peptides 103:31–39. doi:10.1016/j.peptides.2018.03.009

Colmers WF, Wahlestedt C. 1993. The Biology of Neuropeptide Y and related peptides. Springer Science & Business Media.

Costa-da-Silva AL, Navarrete FR, Salvador FS, Karina-Costa M, Ioshino RS, Azevedo DS, Rocha DR, Romano CM, Capurro ML. 2013. Glytube: A Conical Tube and Parafilm M-Based Method as a Simplified Device to Artificially Blood-Feed the Dengue Vector Mosquito, Aedes aegypti. PLoS One 8:e53816. doi:10.1371/journal.pone.0053816

Davis EE. 1984. Regulation of sensitivity in the peripheral chemoreceptor systems for host-seeking behavior by a haemolymph-borne factor in Aedes aegypti. J Insect Physiol 30:179–183. doi:10.1016/0022-1910(84)90124-0.

De Bono M, Bargmann CI. 1998. Natural variation in a neuropeptide Y receptor homolog modifies social behavior and food response in C. elegans. Cell 94:679–689. doi:10.1016/S0092-8674(00)81609-8

Duthaler U, Weber M, Hofer L, Chaccour C, Maia M, Müller P, Kraähenbuähl S, Hammann F. 2021. The pharmacokinetics and drug-drug interactions of ivermectin in Aedes aegypti mosquitoes. PLoS Pathog 17:1–18. doi:10.1371/journal.ppat.1009382

Duvall LB, Ramos-Espiritu L, Barsoum KE, Glickman JF, Vosshall LB. 2019. Small-Molecule Agonists of Ae. aegypti Neuropeptide Y Receptor Block Mosquito Biting. Cell 176:687–701.e5. doi:10.1016/j.cell.2018.12.004

Fiorenzano JM, Koehler PG, Xue R De. 2017. Attractive toxic sugar bait (ATSB) for control of mosquitoes and its impact on non-target organisms: A review. Int J Environ Res Public Health 14. doi:10.3390/ijerph14040398

Galun R. 1963. Feeding Response in Aedes aegypti: Stimulation by Adenosine Triphosphate. Science (80-) 142:1674–1675. doi:10.1126/science.142.3600.1674

Garczynski SF, Crim JW, Brown MR. 2005. Characterization of neuropeptide F and its receptor from the African malaria mosquito, Anopheles gambiae. Peptides 26:99–107. doi:10.1016/j.peptides.2004.07.014

González-Morales MA, Thomson AE, Yeatts J, Enomoto H, Haija A, Santangelo RG, Petritz OA, Crespo R, Schal C, Baynes R. 2023. Pharmacokinetics of fluralaner as a systemic drug to control infestations of the common bed bug, Cimex lectularius, in poultry facilities. Parasites and Vectors 16:1–8. doi:10.1186/s13071-023-05962-3

Gulia-Nuss M, Nuss AB, Meyer JM, Sonenshine DE, Roe RM, Waterhouse RM, Sattelle DB, De La Fuente J, Ribeiro JM, Megy K, Thimmapuram J, Miller JR, Walenz BP, Koren S, Hostetler JB, Thiagarajan M, Joardar VS, Hannick LI, Bidwell S, Hammond MP, Young S, Zeng Q, Abrudan JL, Almeida FC, Ayllón N, Bhide K, Bissinger BW, Bonzon-Kulichenko E, Buckingham SD, Caffrey DR, Caimano MJ, Croset V, Driscoll T, Gilbert D, Gillespie JJ, Giraldo-Calderón GI, Grabowski JM, Jiang D, Khalil SMS, Kim D, Kocan KM, Koči J, Kuhn RJ, Kurtti TJ, Lees K, Lang EG, Kennedy RC, Kwon H, Perera R, Qi Y, Radolf JD, Sakamoto JM, Sánchez-Gracia A, Severo MS, Silverman N, Šimo L, Tojo M, Tornador C, Van Zee JP, Vázquez J, Vieira FG, Villar M, Wespiser AR, Yang Y, Zhu J, Arensburger P, Pietrantonio P V., Barker SC, Shao R, Zdobnov EM, Hauser F, Grimmelikhuijzen CJP, Park Y, Rozas J, Benton R, Pedra JHF, Nelson DR, Unger MF, Tubio JMC, Tu Z, Robertson HM, Shumway M, Sutton G, Wortman JR, Lawson D, Wikel SK, Nene VM, Fraser CM, Collins FH, Birren B, Nelson KE, Caler E, Hill CA. 2016. Genomic insights into the Ixodes scapularis tick vector of Lyme disease. Nat Commun 7. doi:10.1038/ncomms10507

Hamamoto H, Tonoike A, Narushima K, Horie R, Sekimizu K. 2009. Silkworm as a model animal to evaluate drug candidate toxicity and metabolism. Comp Biochem Physiol - C Toxicol Pharmacol 149:334–339. doi:10.1016/j.cbpc.2008.08.008

Hammond A, Galizi R, Kyrou K, Simoni A, Siniscalchi C, Katsanos D, Gribble M, Baker D, Marois E, Russell S, Burt A, Windbichler N, Crisanti A, Nolan T. 2016. A CRISPR-Cas9 gene drive system targeting female reproduction in the malaria mosquito vector Anopheles gambiae. Nat Biotechnol 34:78–83. doi:10.1038/nbt.3439

Harder E, Damm W, Maple J, Wu C, Reboul M, Xiang JY, Wang L, Lupyan D, Dahlgren MK, Knight JL, Kaus JW, Cerutti DS, Krilov G, Jorgensen WL, Abel R, Friesner RA. 2016. OPLS3: A Force Field Providing Broad Coverage of Drug-like Small Molecules and Proteins. J Chem Theory Comput 12. doi:10.1021/acs.jctc.5b00864

He Y, Xu X, Qiu J, Yin W, Sima Y, Xu S. 2020. Bombyx mori used as a fast detection model of liver melanization after a clinical drug – Acetaminophen exposure. J Asia Pac Entomol 23:177–185. doi:10.1016/j.aspen.2019.11.009

Inui A. 1999. Feeding and body-weight regulation by hypothalamic neuropeptides - Mediation of the actions of leptin. Trends Neurosci 22:62–67. doi:10.1016/S0166-2236(98)01292-2

Judson CL. 1967. Feeding and oviposition behavior in the mosquito Aedes aegypti (L.).I. Preliminary studies of physiological control mechanisms. Biol Bull 133:369–377. doi:10.2307/1539832

Khalak Y, Tresadern G, Aldeghi M, Baumann HM, Mobley DL, de Groot BL, Gapsys V. 2021. Alchemical absolute protein-ligand binding free energies for drug design. Chem Sci 12:13958–13971. doi:10.1039/d1sc03472c

Klowden MJ. 1981. Initiation and Termination of Host-Seeking 27:799–803. doi:10.1016/0022-1910(81)90071-8.

Klowden MJ, Lea A. 1979a. Humoral inhibition of host seeking in Aedes aegypti during oocyte maturation 25. doi:10.1016/0022-1910(79)90048-9

Klowden MJ, Lea AO. 1979b. Abdominal distention terminates subsequent host-seeking behaviour of Aedes aegypti following a blood meal. J Insect Physiol 25:583–585. doi:10.1016/0022-1910(79)90073-8

Kraemer MU, Sinka ME, Duda KA, Mylne AQ, Shearer FM, Barker CM, Moore CG, Carvalho RG, Coelho GE, Van Bortel W, Hendrickx G, Schaffner F, Elyazar IR, Teng H-J, Brady OJ, Messina JP, Pigott DM, Scott TW, Smith DL, William Wint G, Golding N, Hay SI. 2015. The global distribution of the arbovirus vectors Aedes aegypti and Ae. albopictus. Elife. doi:10.7554/eLife.08347.001

Krockel U, Rose A, Eiras AE, Geier M. 2006. New tools for surveillance of adult yellow fever mosquitoes: comparison of trap catches with human landing rates in an urban environment. J Am Mosq Control Assoc 22:229–38. doi:10.2987/8756-971X(2006)22[229:NTFSOA]2.0.CO;2

Kuenzel WJ, Douglass LW, Davison BA. 1987. Robust feeding following central administration of neuropeptide Y or peptide YY in chicks, Gallus domesticus. Peptides 8:823–828. doi:10.1016/0196-9781(87)90066-0

Lees RS, Gilles JR, Hendrichs J, Vreysen MJ, Bourtzis K. 2015. Back to the future: the sterile insect technique against mosquito disease vectors. Curr Opin Insect Sci 10:156–162. doi:10.1016/j.cois.2015.05.011

Liesch J, Bellani LL, Vosshall LB. 2013. Functional and Genetic Characterization of Neuropeptide Y-Like Receptors in Aedes aegypti. PLoS Negl Trop Dis 7:e2486. doi:10.1371/journal.pntd.0002486

Liu N. 2015. Insecticide resistance in mosquitoes: Impact, mechanisms, and research directions. Annu Rev Entomol 60:537–559. doi:10.1146/annurev-ento-010814-020828

Macias VM, Ohm JR, Rasgon JL. 2017. Gene drive for mosquito control: Where did it come from and where are we headed? Int J Environ Res Public Health 14. doi:10.3390/ijerph14091006

Moyes CL, Vontas J, Martins AJ, Ng LC, Koou SY, Dusfour I, Raghavendra K, Pinto J, Corbel V, David JP, Weetman D. 2021. Correction to: Contemporary status of insecticide resistance in the major aedes vectors of arboviruses infecting humans (PLoS Negl Trop Dis). PLoS Negl Trop Dis 15:1–2. doi:10.1371/journal.pntd.0009084

Mukabana WR, Mweresa CK, Otieno B, Omusula P, Smallegange RC, van Loon JJA, Takken W. 2012. A Novel Synthetic Odorant Blend for Trapping of Malaria and Other African Mosquito Species. J Chem Ecol 38:235–244. doi:10.1007/s10886-012-0088-8

Okumu FO, Killeen GF, Ogoma S, Biswaro L, Smallegange RC, Mbeyela E, Titus E, Munk C, Ngonyani H, Takken W, Mshinda H, Mukabana WR, Moore SJ. 2010. Development and field evaluation of a synthetic mosquito lure that is more attractive than humans. PLoS One 5. doi:10.1371/journal.pone.0008951

Parihar K, Telang M, Ovhal A. 2020. A patent review on strategies for biological control of mosquito vector. World J Microbiol Biotechnol 36:1–23. doi:10.1007/s11274-020-02960-w

Qiu H, Jun HW, Tao J. 1997. Pharmacokinetics of insect repellent N,N-diethyl-m-toluamide in beagle dogs following intravenous and topical routes of administration. J Pharm Sci 86:514–516. doi:10.1021/js960283m

Ryan SJ, Carlson CJ, Mordecai EA, Johnson LR. 2019. Global expansion and redistribution of Aedes -borne virus transmission risk with climate change. PLoS Negl Trop Dis 13:1–20. doi: 10.1371/journal.pntd.0007213

Savino F, Ricciardi R, Iodice A, Benelli G, Conte G, Lucchi A, Ladurner E, Cosci F. 2018. Eco-friendly pheromone dispensers—a green route to manage the European grapevine moth? Environ Sci Pollut Res 25:9426–9442. doi:10.1007/s11356-018-1248-3

Su X, Guo Y, Deng J, Xu J, Zhou G, Zhou T, Li Y, Zhong D, Kong L, Wang X, Liu M, Wu K, Yan G, Chen XG. 2019. Fast emerging insecticide resistance in Aedes albopictus in Guangzhou, China: Alarm to the dengue epidemic. PLoS Negl Trop Dis 13:1–15. doi:10.1371/journal.pntd.0007665

Weaver SC, Costa F, Garcia-Blanco MA, Ko AI, Ribeiro GS, Saade G, Shi PY, Vasilakis N. 2013. Zika virus: History, emergence, biology, and prospects for control. PLoS One 13:1–15. doi:10.1155/2017/3196924

Witmer GW, Raymond-Whish S, Moulton RS, Pyzyna BR, Calloway EM, Dyer CA, Mayer LP, Hoyer PB. 2017. Compromised Fertility in Free Feeding of Wild-Caught Norway Rats (Rattus Norvegicus) With a Liquid Bait Containing 4-Vinylcyclohexene Diepoxide and Triptolide. J Zoo Wildl Med 48:80–90. doi:10.1638/2015-0250.1

World Health Organization. 2012. Global Plan for Insecticide Resistance Management in Malaria Vectors. Geneva, Swizterland. doi:WHO REFERENCE NUMBER: WHO/HTM/GMP/2012.5

Wu Q, Wen T, Lee G, Park JH, Cai HN, Shen P. 2003. Developmental control of foraging and social behavior by the Drosophila neuropeptide Y-like system. Neuron 39:147–161. doi:10.1016/S0896-6273(03)00396-9

Zhang J, Yang J, Jang R, Zhang Y. 2015. GPCR-I-TASSER: A Hybrid Approach to G Protein-Coupled Receptor Structure Modeling and the Application to the Human Genome. Structure 23:1538–1549. doi:10.1016/j.str.2015.06.007

