## Supplemental methods for synthesis reactions for "Next Generation Neuropeptide Y Receptor Small Molecule Agonists Inhibit Mosquito Biting Behavior"

Synthetic methods for analog preparation

Unless otherwise noted, the following pertain to the synthetic methods: all reactions are magnetically stirred; typical solvents (ethyl acetate, hexanes, dichloromethane, and methanol) are Fisher Optima grade; “concentrated to dryness” or “removal of the solvent” means evaporating the solvent from a solution or mixture using a rotary evaporator; flash chromatography is carried out on an Isco, Analogix, or Biotage automated chromatography system using a commercially available cartridge as the column. Columns are usually filled with silica gel as the stationary phase; preparative HPLC (or prep-HPLC) is carried out with commercial columns in a reverse phase manner (the stationary phase is hydrophobic). Typical solvent mixtures include A (water) and B (organic i.e. acetonitrile, methanol, etc.). Additives can also be used in the solvent mixture such as HCl, NH_4_HCO_3_, and formic acid.

Preparation of TDI-012610

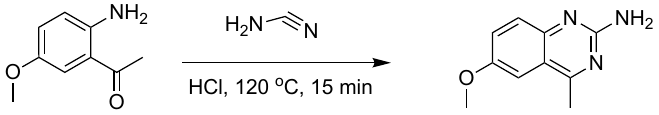

A mixture of 1-(2-amino-5-methoxy-phenyl)ethanone (0.200 g, 1.21 mmol, 1 eq), HCl (0.3 mL, 1M aq.) in cyanamide (2.04 g, 24.21 mmol, 2.04 mL, 50% wt in H_2_O) was stirred at 120 °C for 15 min. The reaction mixture was cooled to 20 °C and diluted with H_2_O (8 mL). The mixture was concentrated under vacuum. The residue was purified by prep-HPLC (HCl condition). [column: Welch Xtimate C18 100 x 25 mm x 3 mm; mobile phase: [water (0.05% HCl)-ACN];B%: 1%-15%,8min] to furnish 6-methoxy-4-methyl-quinazolin-2-amine (0.045 g, 0.23 mmol, 19%) as a yellow solid. LCMS (M+H)^+^ = 190.0; 1H NMR: (400 MHz, DMSO-d6) δ 7.69-7.59 (m, 2H), 7.56 (d, *J* = 1.8 Hz, 1H), 3.92 (s, 3H), 2.89 (s, 3H).

Preparation of TDI-014179

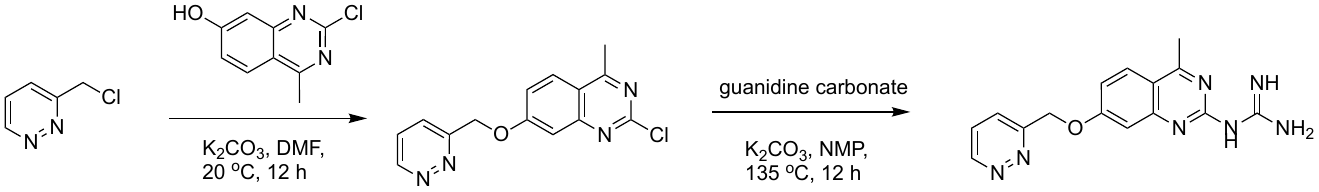

To a solution of 3-(chloromethyl)pyridazine (130 mg, 1.01 mmol, 1 eq) in DMF (2 mL) was added K_2_CO_3_ (419 mg, 3.03 mmol, 3 eq) and 2-chloro-4-methyl-quinazolin-7-ol (200 mg, 1.03 mmol, 1.02 eq) . The mixture was stirred at 25 °C for 12 h. The mixture was poured into H_2_O (10 mL), and the mixture was extracted with ethyl acetate ( 5 mL x 3). The combined organic layers were washed with brine (10 mL), dried over Na_2_SO_4_, and filtered. The filtrate was concentrated under reduced pressure to give a residue. The residue was purified by column chromatography (SiO_2_, petroleum ether/ethyl acetate = 9/1 to 0/1) to furnish 2-chloro-4-methyl-7-(pyridazin-3-ylmethoxy)quinazoline (100 mg, crude) as a white solid.

To a solution of 2-chloro-4-methyl-7-(pyridazin-3-ylmethoxy)quinazoline (80 mg, 0.28 mmol, 1 eq) in NMP (2 mL) was added K_2_CO_3_ (116 mg, 0.84 mmol, 3 eq) and guanidine carbonate (55 mg, 0.31 mmol, 1.1 eq) .The mixture was stirred at 135 °C for 12 hr . The reaction mixture was concentrated under reduced pressure. The residue was purified by preparative-HPLC (HCl conditions). column: Phenomenex luna C18 80 x 40mm x 3 mm; mobile phase: [water(HCl)-ACN];B%: 5%-30%,7min to furnish 1-[4-methyl-7-(pyridazin-3-ylmethoxy)quinazolin-2-yl]guanidine (2 mg, 0.006 mmol, HCl salt) as a solid. LCMS (M+H)^+^ = 310.0; 1H NMR: (400 MHz, DMSO-d6) δ 11.07 (br s, 1H), 9.27 (dd, J = 1.5, 4.9 Hz, 1H), 8.71-8.36 (m, 3H), 8.20 (d, J = 9.1 Hz, 1H), 7.93 (dd, J = 1.4, 8.5 Hz, 1H), 7.82 (dd, J = 5.0, 8.5 Hz, 1H), 7.60 (d, J = 2.5 Hz, 1H), 7.35 (dd, J = 2.5, 9.1 Hz, 1H), 5.61 (s, 2H), 2.85 (s, 3H).

Preparation of TDI-014186

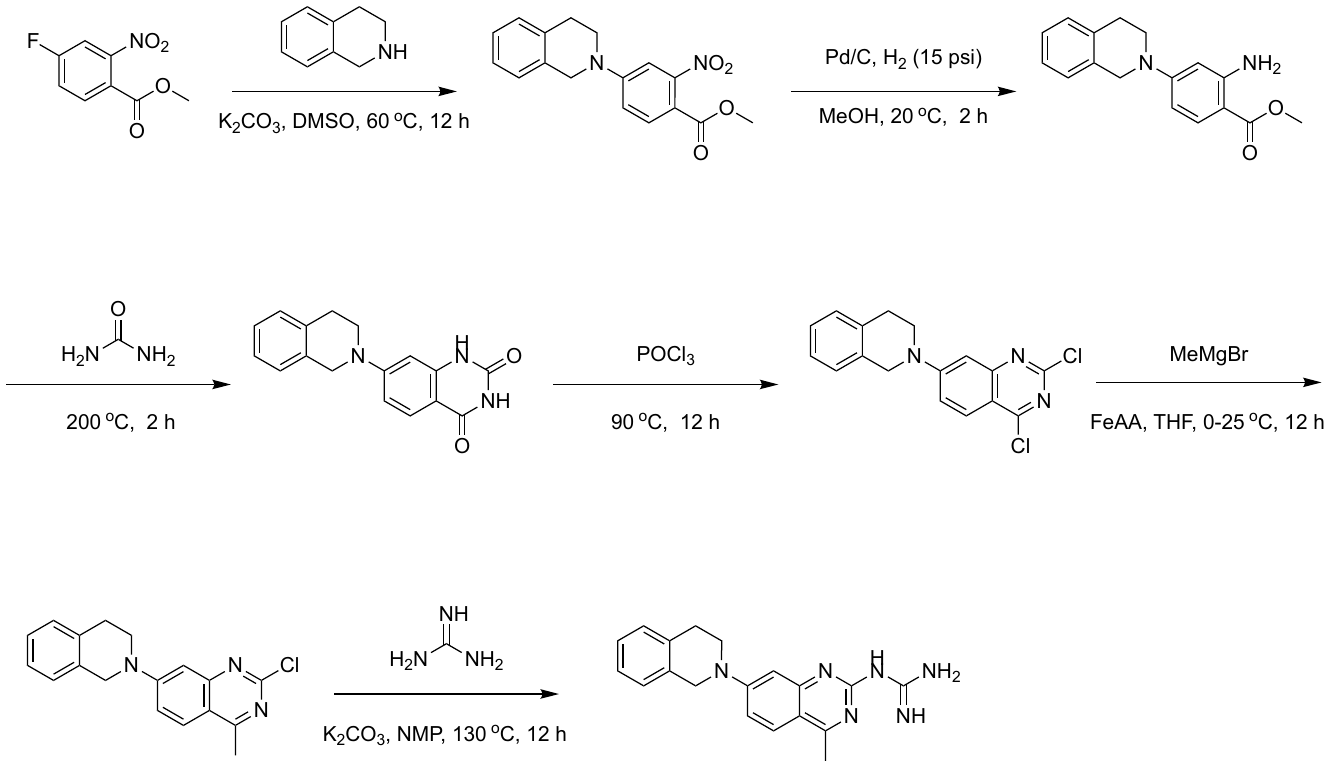

A mixture of methyl 4-fluoro-2-nitro-benzoate (4.00 g, 20.1 mmol, 1 *eq*), 1,2,3,4- tetrahydroisoquinoline (3.21 g, 24.1 mmol, 3.03 mL, 1.2 *eq*) and K_2_CO_3_ (5.83 g, 42.2 mmol, 2.1 *eq*) in DMSO (20 mL) was degassed and purged with N_2_ (3X). The mixture was stirred at 60 °C for 12 h under a N_2_ atmosphere. The mixture was poured into water (20 mL), and the mixture was extracted with ethyl acetate (15 mL x 3). The combined organic phase was washed with brine (15 mL x 1), dried with anhydrous Na_2_SO_4_, and filtered. The filtrate was concentrated to furnish methyl 4-(3,4-dihydro-1H-isoquinolin-2-yl)-2-nitro-benzoate (3.7 g, crude) as a yellow solid.

To a solution of methyl 4-(3,4-dihydro-1H-isoquinolin-2-yl)-2-nitro-benzoate (3.70 g, 11.9 mmol, 1 eq) in MeOH (40 mL) was added Pd/C (3.8 g, 10 wt% on carbon) under a N_2_ atmosphere. The suspension was degassed and purged with H_2_ (3X). The mixture was stirred under H_2_ (15 Psi ) at 20 °C for 2 h. The mixture was filtered, and the filtrate was concentrated under reduced pressure to furnish methyl 2-amino-4-(3,4-dihydro-1H-isoquinolin-2-yl)benzoate (2.1 g, crude) as a yellow oil.

A mixture of methyl 2-amino-4-(3,4-dihydro-1H-isoquinolin-2-yl)benzoate (2.10 g, 7.44 mmol, 1 eq), urea (5.36 g, 89.3 mmol, 12 eq) was degassed and purged with N_2_ for (3X). The mixture was stirred at 200 °C for 2 h under a N_2_ atmosphere. The resulting mixture was treated with water (50 mL) at 100 °C. The resulting precipitate was filtered and washed with EtOH/H_2_O(1:1, 100 mL) to furnish 7-(3,4-dihydro-1H-isoquinolin-2-yl)-1H-quinazoline-2,4-dione (2.5 g, crude) as a yellow solid.

A mixture of 7-(3,4-dihydro-1H-isoquinolin-2-yl)-1H-quinazoline-2,4-dione (2.50 g, 8.52 mmol, 1 *eq*) in POCl_3_ (30 mL) was degassed and purged with N_2_ (3X). The mixture was stirred at 90 °C for 12 h under a N_2_ atmosphere. The mixture was quenched with aq. Na_2_CO_3_ (30 mL), and the mixture was extracted with EtOAc( 20 mL x 3). The combined organic layers were washed with brine (30 mL), dried overNa_2_SO_4_, and filtered. The filtrate was concentrated under reduced pressure to give a residue. The residue was purified by flash silica gel chromatography (ISCO®; 25 g SepaFlash® Silica Flash Column, eluent of 0 to 50% ethyl acetate/petroleum ether gradient at 75 mL/min) to furnish 2,4-dichloro-7-(3,4-dihydro-1H-isoquinolin-2-yl)quinazoline (600 mg, 1.45 mmol, 17 % yield) as a yellow solid. 1H NMR: (400MHz, DMSO-d6) δ 8.02 (d, J = 9.5 Hz, 1H), 7.69 (dd, J = 2.4, 9.6 Hz, 1H), 7.33-7.29 (m, 1H), 7.26-7.21 (m, 3H), 7.15-7.13 (m, 1H), 4.73 (s, 2H), 3.81 (t, J = 5.9 Hz, 2H), 3.02-2.98 (m, 2H).

To a solution of 2,4-dichloro-7-(3,4-dihydro-1H-isoquinolin-2-yl)quinazoline(214 mg, 0.648 mmol, 1 eq) in THF (10 mL) was added tris[(Z)-1-methyl-3-oxo-but-1-enoxy]iron (69 mg, 0.19 mmol, 0.3 eq). After the addition, MeMgBr (3 M, 0.320 mL, 1.5 eq) was added dropwise at 0 °C. The resulting mixture was stirred at 25 °C for 12 h. The residue was diluted with NH_4_Cl (50 mL), and the mixture was extracted with ethyl acetate (50 mL x 3). The combined organic layer was washed with NaCl (20 mL x 3), dried over Na_2_SO_4_, and filtered. The filtrate was concentrated under reduced pressure. The residue was purified by flash silica gel chromatography (ISCO®; 12 g SepaFlash® Silica Flash Column, eluent of 0 to 50% ethyl acetate/petroleum ether gradient at 75 mL/min) to furnish 2-chloro-7-(3,4-dihydro-1H- isoquinolin-2-yl)-4-methyl-quinazoline (70 mg, 0.14 mmol, 21 % yield) as a yellow oil.

A mixture of 2-chloro-7-(3,4-dihydro-1H-isoquinolin-2-yl)-4-methyl-quinazoline (70 mg, 0.23 mmol, 1 eq), guanidine (20 mg, 0.34 mmol, 1.5 eq) and K_2_CO_3_ (63 mg, 0.45 mmol, 2 eq) in NMP (1 mL) was degassed and purged with N_2_ (3X). The mixture was stirred at 130 °C for 12 h under a N_2_ atmosphere. The mixture was filtered, and the filtrate was concentrated under reduced pressure. The residue was purified by prep-HPLC (neutral condition). (column: Waters Xbridge BEH C18 100 x 30mm x 10 mm; mobile phase: [water( NH_4_HCO_3_)-ACN];B%: 25%-55%, 8 min) to furnish 1-[7-(3,4-dihydro-1H-isoquinolin-2-yl)-4-methyl-quinazolin-2-yl]guanidine (9 mg, 22 mmol, 10 % yield) as a white solid. LCMS (M+H)^+^ = 310.0; 1H NMR: (400 MHz, DMSO-d6) δ 11.07 (br s, 1H), 9.27 (dd, J = 1.5, 4.9 Hz, 1H), 8.71-8.36 (m, 3H), 8.20 (d, J = 9.1 Hz, 1H), 7.93 (dd, J = 1.4, 8.5 Hz, 1H), 7.82 (dd, J = 5.0, 8.5 Hz, 1H), 7.60 (d, J = 2.5 Hz, 1H), 7.35 (dd, J = 2.5, 9.1 Hz, 1H), 5.61 (s, 2H), 2.85 (s, 3H).

Preparation of TDI-014172

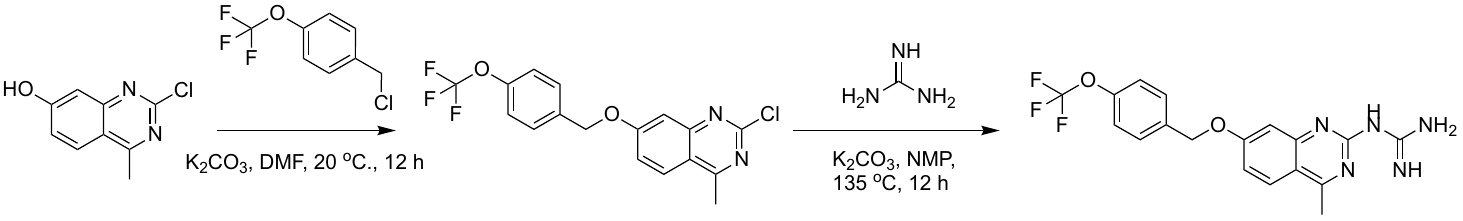

A mixture of 2-chloro-4-methyl-quinazolin-7-ol (0.100 g, 0.514 mmol, 1 eq), 1-(chloromethyl)-4- (trifluoromethoxy)benzene (162 mg, 0.771 mmol, 1.5 eq), K_2_CO_3_ (213 mg, 1.54 mmol, 3 eq) in DMF (1 mL) was degassed and purged with N_2_ (3X). The mixture was stirred at 20 °C for 12 h under a N_2_ atmosphere. The mixture was quenched with H_2_O (10 mL), and the mixture was extracted with ethyl acetate (10 mL x 3). The combined organic layers were concentrated under reduced pressure to give a residue. The residue was purified by column chromatography (SiO_2_, petroleum ether/ethyl acetate = 5/1) to furnish 2-chloro-4-methyl-7-((4-(trifluoromethoxy)benzyl)oxy)quinazoline (40 mg) as a white solid.

A mixture of 2-chloro-4-methyl-7-((4-(trifluoromethoxy)benzyl)oxy)quinazoline (40 mg, 0.11 mmol, 1 eq), guanidine carbonate (65 mg, 0.36 mmol, 3.3 eq), K_2_CO_3_ (75 mg, 0.54 mmol, 5 eq) in NMP (1 mL) was degassed and purged with N_2_ (3X). The mixture was stirred at 135 °C for 12 h under a N_2_ atmosphere. The mixture was filtered and concentrated under reduced pressure to give a residue. The residue was purified by prep-HPLC (HCl conditions, column: Phenomenex Luna 80 x 30mm x 3 mm; mobile phase: water(HCl)-ACN];B%: 20%-60%, 8 min) to furnish 1-(4-methyl-7-((4-(trifluoromethoxy) benzyl)oxy)quinazolin-2-yl)guanidine (22 mg, 0.055 mmol, 51 % yield) as a white solid. LCMS (M+H)^+^ = 392.0; 1H NMR: (400MHz, DMSO-d6) δ 11.03 (s, 1H), 8.68-8.37 (m, 3H), 8.18 (d, J = 9.0 Hz, 1H), 7.66 (d, J = 8.6 Hz, 2H), 7.53 (d, J = 2.4 Hz, 1H), 7.44 (d, J = 8.3 Hz, 2H), 7.31 (dd, J = 2.5, 9.1 Hz, 1H), 5.34 (s, 2H), 2.84 (s, 3H).

Preparation of TDI-014183

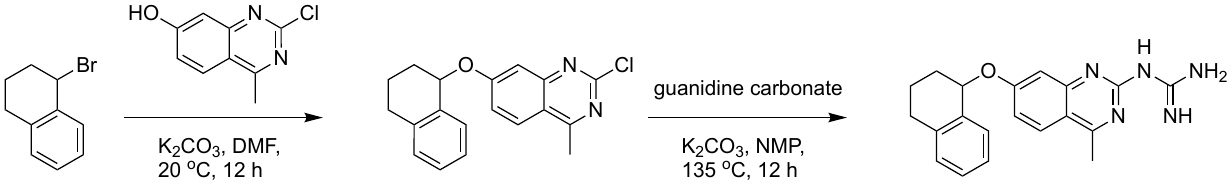

To a solution of 2-chloro-4-methyl-quinazolin-7-ol (200 mg, 1.03 mmol, 1 eq) in DMF (4 mL) was added K_2_CO_3_ (426 mg, 3.08 mmol, 3 eq) and 1-bromotetralin (220 mg, 1.04 mmol, 1.01 eq) . The mixture was stirred at 20 °C for 12 h. The reaction mixture was concentrated under reduced pressure. The mixture was poured into H_2_O (10 mL), and then the mixture was extracted with ethyl acetate (5 mL x 3). The combined organic layers were washed with brine (10 mL), dried over Na_2_SO_4_, and filtered. The filtrate was concentrated under reduced pressure to give a residue. The residue was purified by column chromatography (SiO_2_, petroleum ether/ethyl acetate = 9/1 to 5/1) to furnish 2-chloro-4-methyl-7-tetralin-1-yloxy-quinazoline (60 mg, crude) as a yellow solid.

To a solution of 2-chloro-4-methyl-7-tetralin-1-yloxy-quinazoline (60 mg, 0.19 mmol, 1 eq) in NMP (1 mL) was added K_2_CO_3_ (128 mg, 0.924 mmol, 5 eq) and guanidine carbonate (37 mg, 0.20 mmol, 1.1 eq) . The mixture was stirred at 135 °C for 12 h. The reaction mixture was concentrated under reduced pressure. The residue was purified by prep-HPLC (neutral condition, column: Phenomenex luna C18 80 x 40 mm x 3 mm; mobile phase: [water(HCl)-ACN];B%: 27%-45%,7min) to furnish 1-(4-methyl-7-tetralin-1-yloxy-quinazolin-2-yl)guanidine (10 mg, 0.025 mmol, 14 %, HCl) as a white solid. LCMS (M+H)^+^ = 348.1; 1H NMR: (400 MHz, DMSO-d6) δ 10.97 (s, 1H), 8.58-8.25 (m, 3H), 8.17 (d, *J* = 9.1 Hz, 1H), 7.66 (d, *J* = 2.4 Hz, 1H), 7.33-7.24 (m, 3H), 7.20 (d, *J* = 7.4 Hz, 2H), 5.78 (t, *J* = 4.3 Hz, 1H), 2.92-2.86 (m, 1H), 2.84 (s, 3H), 2.80-2.74 (m, 1H), 2.13-2.07 (m, 2H), 1.94-1.87 (m, 1H), 1.85-1.78 (m, 1H).

Preparation of TDI-013758

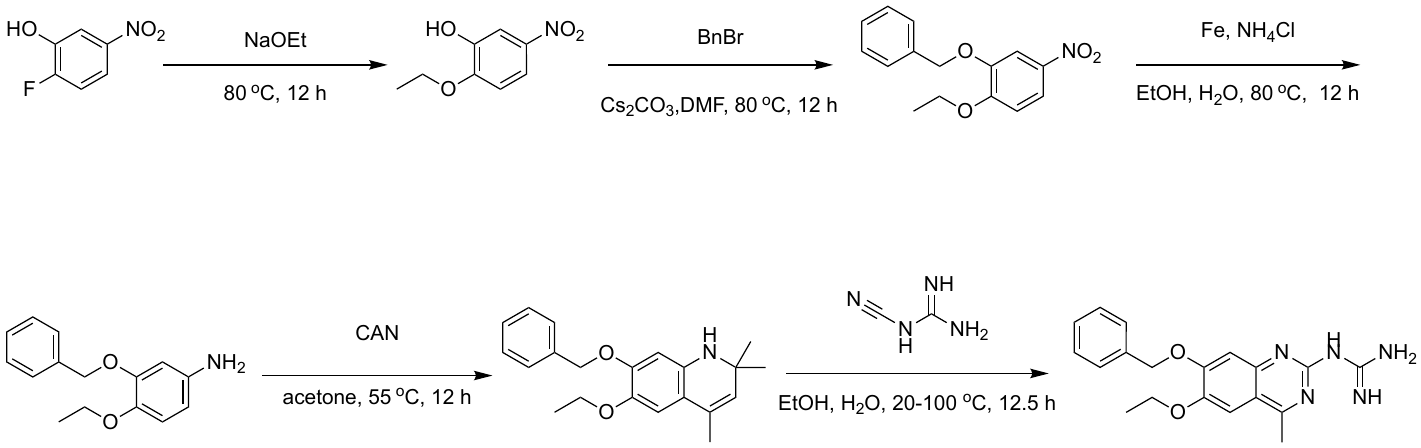

A mixture of 2-fluoro-5-nitro-phenol (3.00 g, 19.1 mmol, 1 *eq*) in NaOEt (39.0, 115 mmol, 20% in ethanol, 6 *eq*) was degassed and purged with N_2_ (3X). The mixture was stirred at 80 °C for 12 h under a N_2_ atmosphere. The mixture was poured into water (100 mL), and the mixture was treated with aqueous HCl (2M, 30 mL) to achieve a final pH of 3. The mixture was extracted with ethyl acetate (150 mL x 3). The combined organic phase was washed with brine (50 mL), dried with anhydrous Na_2_SO_4_, and filtered. The filtrate was concentrated. The residue was purified by flash silica gel chromatography (ISCO®; 12 g SepaFlash® silica flash column, eluent of 0 to 50% ethyl acetate/petroleum ether gradient @ 75 mL/min) to furnish 2-ethoxy-5-nitro-phenol (2.5 g, crude) as a brown solid.

To a solution of 2-ethoxy-5-nitro-phenol (2.50 g, 13.7 mmol, 1 *eq*) in DMF (30 mL) was added Cs_2_CO_3_ (8.89 g, 27.3 mmol, 2 *eq*) and benzyl bromide (2.80 g, 16.4 mmol, 1.95 mL, 1.2 *eq*). The mixture was stirred at 80 °C for 12 h. The mixture was poured into water (50 mL), and the formed solid was collected and dried under reduced pressure to furnish 2-benzyloxy-1-ethoxy- 4-nitro-benzene (2.2 g, crude) as a yellow solid.

A mixture of 2-benzyloxy-1-ethoxy-4-nitro-benzene (2.20 g, 8.05 mmol, 1 *eq*), Fe (2.25 g, 40.3 mmol, 5 *eq*) and NH_4_Cl (2.15 g, 40.3 mmol, 5 *eq*) in EtOH (50 mL) and H_2_O (10 mL) was degassed and purged with N_2_ (3X). The mixture was stirred at 80 °C for 12 h under a N_2_ atmosphere. The mixture was filtered, and the filtrate was concentrated under reduced pressure to furnish 3-benzyloxy-4-ethoxy-aniline (1.8 g, crude) as a brown solid.

To a solution of 3-benzyloxy-4-ethoxy-aniline (1.80 g, 7.40 mmol, 1 *eq*) in acetone (5 mL) was added ceric ammonium nitrate (1.01 g, 1.85 mmol, 0.25 *eq*). The mixture was stirred at 55 °C for 12 h. The reaction mixture was concentrated under reduced pressure to remove solvent. The residue was purified by flash silica gel chromatography (ISCO®; 40 g SepaFlash® silica flash column, eluent of 0 to 20% ethyl acetate/petroleum ether gradient at 100 mL/min) to furnish 7-benzyloxy-6-ethoxy-2,2,4-trimethyl-1H-quinoline (860 mg, 2.39 mmol, 32 %) as a yellow solid.

After stirring a solution of 7-benzyloxy-6-ethoxy-2,2,4-trimethyl-1H-quinoline (280 mg, 0.778 mmol, 1 *eq*, HCl) in EtOH (3 mL) and H_2_O (3 mL) for 30 min, 1-cyanoguanidine (79 mg, 0.93 mmol, 1.2 *eq*) was added. The resulting mixture was stirred at 100 °C for 12 h. The reaction mixture was concentrated under reduced pressure to remove solvent. The mixture was poured into DMSO (4 mL). The mixture was filtered, and the filtrate was concentrated under reduced pressure to give a residue. The residue was purified by prep-HPLC (HCl condition). (column: Phenomenex Luna 80 x 30mm x 3 mm; mobile phase: [water(0.04%HCl)-ACN];B%: 25%-55%,8min) to furnish 1-(7-benzyloxy -6-ethoxy-4-methyl-quinazolin-2-yl)guanidine (23 mg, 0.065 mmol, 8 %) as a white solid. LCMS (M+H)^+^ = 352.0; ^1^H NMR: (400MHz, METHANOL-d4) δ 7.50 (d, J = 7.23 Hz, 2H) 7.43-7.38 (m, 2H) 7.37-7.31 (m, 3H) 5.27 (s, 2H), 4.21 (q, J = 6.87 Hz, 2H), 2.81 (s, 3H), 1.49 (t, J = 7.02 Hz, 3H.

Preparation of TDI-014167

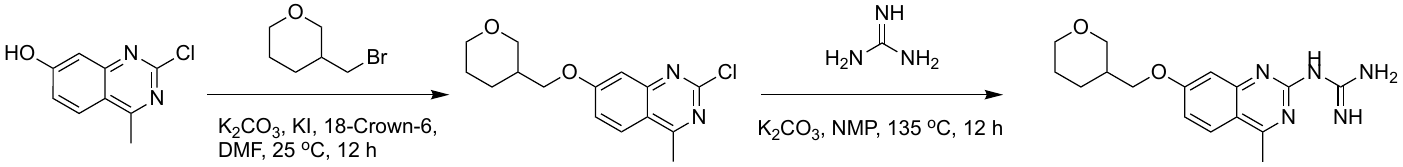

A mixture of 2-chloro-4-methyl-quinazolin-7-ol (100 mg, 0.514 mmol, 1 *eq*), 3- (bromomethyl)tetrahydropyran (138 mg, 0.771 mmol, 1.5 *eq*), K_2_CO_3_ (213 mg, 1.54 mmol, 3 *eq*), KI (9 mg, 0.051 mmol, 0.1 *eq*) and 1,4,7,10,13,16-hexaoxacyclooctadecane (14 mg, 0.051 mmol, 0.1 *eq*) in DMF (1 mL) was degassed and purged with N_2_ (3X). The mixture was stirred at 25 °C for 12 h under a N_2_ atmosphere. The mixture was poured into water (20 mL), and the mixture was extracted with ethyl acetate (15 mL x 3). The combined organic phase was washed with brine (15 mL), dried with anhydrous Na_2_SO_4_, and filtered. The filtrate was concentrated to furnish 2-chloro-4-methyl-7-(tetrahydropyran-3-ylmethoxy) quinazoline (130 mg, crude) as a yellow solid.

A mixture of 2-chloro-4-methyl-7-(tetrahydropyran-3-ylmethoxy)quinazoline (100 mg, 0.341 mmol, 1 *eq*), guanidine carbonate (123 mg, 0.683 mmol, 2 *eq*) and K_2_CO_3_ (94 mg, 0.68 mmol, 2 *eq*) in NMP (1 mL) was degassed and purged with N_2_ (3X). The mixture was stirred at 135 °C for 12 h under a N_2_ atmosphere. The mixture was poured into water (10 mL), and the mixture was extracted with ethyl acetate (15 mL x 3). The combined organic phase was washed with brine (15 mL), dried with anhydrous Na_2_SO_4_, and filtered. The filtrate was concentrated. The residue was purified by prep-HPLC (HCl condition). (column: Phenomenex luna C18 80 x 40mm x 3 mm; mobile phase: [water(HCl)-ACN];B%: 24%-44%,7min) to furnish 1-[4-methyl-7- (tetrahydropyran-3-ylmethoxy)quinazolin-2-yl]guanidine (14 mg) as a white solid. LCMS (M+H)^+^ = 316.0; ^1^H NMR: (400MHz, METHANOL-d4) δ 8.11 (d, J = 9.13 Hz, 1H), 7.31 (d, J = 2.38 Hz, 1H), 7.22 (dd, J = 9.13, 2.50 Hz, 1H), 4.10-4.02 (m, 2H), 3.86 (dt, J = 11.16, 3.80 Hz, 1H), 3.53-3.41 (m, 2H), 2.87-2.84 (m, 3H), 2.41 (t, J = 8.07 Hz, 1H), 2.24-2.14 (m, 1H), 2.08-1.94 (m, 1H), 1.79-1.60 (m, 2H), 1.60-1.48 (m, 1H).

Preparation of TDI-014176

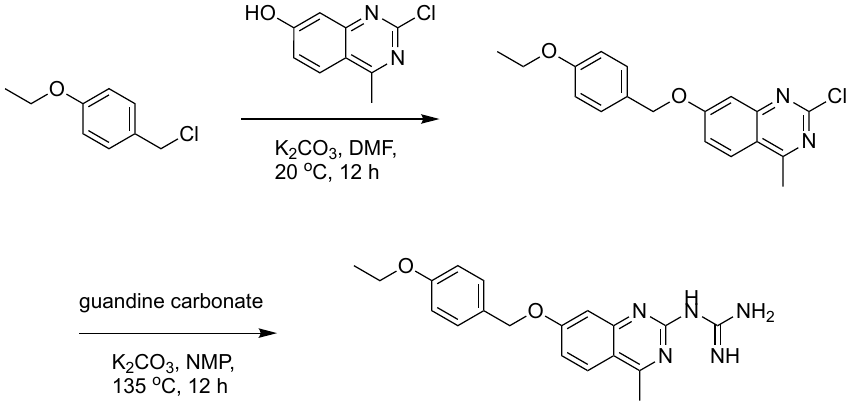

To a solution of 2-chloro-4-methyl-quinazolin-7-ol (200 mg, 1.03 mmol, 1 eq) in DMF (4 mL) was added K_2_CO_3_ (426 mg, 3.08 mmol, 3 eq) and 1-(chloromethyl)-4-ethoxy-benzene (180 mg, 1.05 mmol, 1.03 eq) . The mixture was stirred at 20 °C for 12 h. The reaction mixture was concentrated under reduced pressure. The mixture was poured into H_2_O (10 mL), and the mixture was extracted with ethyl acetate (5 mL x 3). The combined organic layers were washed with brine (10 mL), dried over Na_2_SO_4_, and filtered. The filtrate was concentrated under reduced pressure to give a residue. The residue was purified by column chromatography (SiO_2_, petroleum ether/ethyl acetate = 9/1 to 5/1) to furnish 2-chloro-7-[(4-ethoxyphenyl)methoxy]-4-methyl-quinazoline (200 mg, crude) as a yellow solid.

To a solution of 2-chloro-7-[(4-ethoxyphenyl)methoxy]-4-methyl-quinazoline (100 mg, 0.304 mmol, 1 eq) in NMP (1 mL) was added K_2_CO_3_ (210 mg, 1.52 mmol, 5 eq) and guanidine carbonate (60 mg, 0.334 mmol, 1.1 eq) . The mixture was stirred at 135 °C for 12 h. The reaction mixture was concentrated under reduced pressure. The residue was purified by prep-HPLC (neutral condition, column: Waters Xbridge BEH C18 100 x 30mm x 10 mm; mobile phase: [water( NH_4_HCO_3_)-ACN];B%: 35%-65%,10min) to furnish 1-[7-[(4-ethoxyphenyl)methoxy]-4-methyl-quinazolin-2-yl]guanidine (3 mg, 0.008 mmol, 3 %) as a white solid. LCMS (M+H)^+^ = 352.1; ^1^H NMR: (400MHz, DMSO-d6) δ 7.85 (d, *J* = 9.0 Hz, 1H), 7.40 (d, *J* = 8.5 Hz, 2H), 7.06 (br d, *J* = 2.4 Hz, 3H), 6.98-6.87 (m, 3H), 5.13 (s, 2H), 4.03 (q, *J* = 7.0 Hz, 2H), 2.63 (s, 3H), 1.32 (t, *J* = 6.9 Hz, 3H).

Preparation of TDI-014181

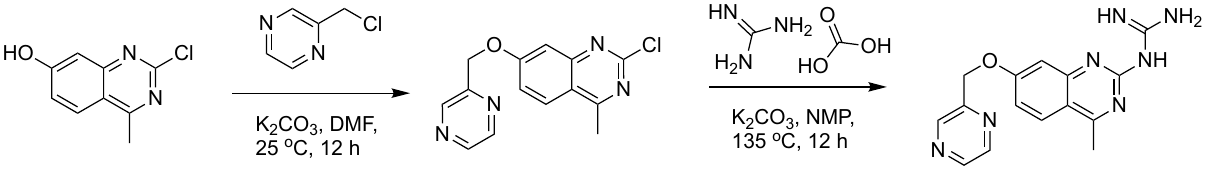

To a solution of 2-chloro-4-methyl-quinazolin-7-ol (130 mg, 0.668 mmol, 1 eq) in DMF (2 mL) was added K_2_CO_3_ (277 mg, 2.00 mmol, 3 eq) and 2-(chloromethyl) pyrazine (130 mg, 1.01 mmol, 1.51 eq). The mixture was stirred at 25 °C for 12 h. The mixture was poured into H_2_O (10 mL), and then the mixture was extracted with ethyl acetate (5 mL x 3). The combined

organic layers were washed with brine (10 mL), dried over Na_2_SO_4_, and filtered. The filtrate was concentrated under reduced pressure to give a residue. The residue was purified by column chromatography (SiO_2_, petroleum ether/ethyl acetate = 9/1 to 1/3) to furnish 2-chloro-4-methyl-7-(pyrazin-2-ylmethoxy) quinazoline (30 mg, crude) as a white solid.

To a solution of 2-chloro-4-methyl-7-(pyrazin-2-ylmethoxy) quinazoline (30 mg, 0.10 mmol, 1 eq) in NMP (1 mL) was added K_2_CO_3_ (43 mg, 0.31 mmol, 3 eq) and guanidine carbonate (21 mg, 0.12 mmol, 1.1 eq). The mixture was stirred at 135 °C for 12 h. The mixture was filtered and concentrated in vacuum. The residue was purified by Prep-HPLC (neutral condition; column: Waters Xbridge BEH C18 100 x 30 mm x 10 mm; mobile phase: [water (NH_4_HCO_3_)-ACN]; B%: 30%-60%, 8min) to furnish 1-[4-methyl-7-(pyrazin-2-ylmethoxy) quinazolin-2-yl] guanidine (1.2 mg, 0.004 mmol, 4%) as a white solid. LCMS (M+H)^+^ = 310.0; ^1^H NMR: (400MHz, DMSO-d6) δ 8.87 (s, 1H), 8.68 (dd, *J* = 1.9, 14.8 Hz, 2H), 7.95 (d, *J* = 8.9 Hz, 1H), 7.53-7.29 (m, 2H), 7.21 (br s, 1H), 7.10-7.04 (m, 1H), 5.40 (s, 2H), 2.68 (s, 3H).

Preparation of TDI-014192

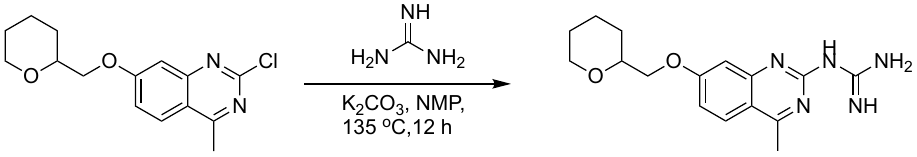

To a solution of 2-chloro-4-methyl-7-(tetrahydropyran-2-ylmethoxy)quinazoline (120 mg, 0.410 mmol, 1 eq) in NMP (2 mL) was added K_2_CO_3_ (170 mg, 1.23 mmol, 3 eq) and guanidine carbonate (81 mg, 0.45 mmol, 1.1eq). The mixture was stirred at 135 °C for 12 h. The reaction mixture was concentrated under reduced pressure. The residue was purified by prep-HPLC (formic acid[FA] conditions, column: Phenomenex Luna C18 150 x 30 mm x 5 mm; mobile phase: [water(FA)-ACN];B%: 1%-40%,8min) to furnish 1-[4-methyl -7-(tetrahydropyran-2-ylmethoxy)quinazolin-2-yl]guanidine (30 mg, 0.090 mmol, 22 %) as a white solid. LCMS (M+H)^+^ = 316.1; ^1^H NMR: (400MHz, DMSO-d6) δ 9.84-8.83 (m, 3H), 8.46 (s, 1H), 8.06 (d, *J* = 9.2 Hz, 1H), 7.37 (d, *J* = 2.3 Hz, 1H), 7.16 (dd, *J* = 2.4, 9.0 Hz, 1H), 4.12-4.04 (m, 2H), 3.90 (br d, *J* = 11.9 Hz, 1H), 3.72-3.66 (m, 1H), 3.42-3.37 (m, 1H), 2.78 (s, 3H), 1.83 (br d, *J* = 8.7 Hz, 1H), 1.67 (br d, *J* = 12.0 Hz, 1H), 1.57-1.43 (m, 3H), 1.41 - 1.28 (m, 1H).

Preparation of TDI-014177

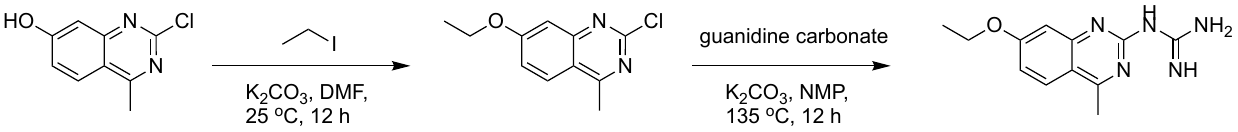

To a solution of 2-chloro-4-methyl-quinazolin-7-ol (200 mg, 1.03 mmol, 1 eq) in DMF (4 mL) was added K_2_CO_3_ (426 mg, 3.08 mmol, 3 eq) and iodoethane (167 mg, 1.07 mmol, 1.04 eq) . The mixture was stirred at 25 °C for 12 h. The reaction mixture was concentrated under reduced pressure. The mixture was poured into H_2_O (10 mL), and the mixture was extracted with ethyl acetate (5 mL x 3). The combined organic layers were washed with brine (10 mL), dried over Na_2_SO_4_, and filtered. The filtrate was concentrated under reduced pressure to give a residue. The residue was purified by column chromatography (SiO_2_, petroleum ether/ethyl acetate = 9/1 to 5/1) to furnish 2-chloro-7-ethoxy-4-methyl-quinazoline (180 mg, crude) as a yellow solid.

To a solution of 2-chloro-7-ethoxy-4-methyl-quinazoline (150 mg, 0.674 mmol, 1 eq) in NMP (3 mL) was added K_2_CO_3_ (466 mg, 3.37 mmol, 5 eq) and guanidine carbonate (134 mg, 0.741 mmol, 1.1 eq) . The mixture was stirred at 135 °C for 12 h. The mixture was filtered and concentrated in vacuum. The solid was washed with ethyl acetate (5 mL) and H_2_O (5 mL) to furnish 1-(7-ethoxy-4-methyl-quinazolin-2-yl)guanidine (54 mg, 0.22 mmol, 32%) as a yellow solid. LCMS (M+H)^+^ = 246.0; ^1^H NMR: (400MHz, DMSO-d6) δ 7.83 (d, *J* = 9.0 Hz, 1H), 7.07 (br s, 2H), 6.92 (d, *J* = 2.4 Hz, 1H), 6.85 (dd, *J* = 2.5, 9.0 Hz, 1H), 4.14 (q, *J* = 6.9 Hz, 2H), 2.62 (s, 3H), 1.38 (t, J = 6.9 Hz, 3H).

Preparation of TDI-014174

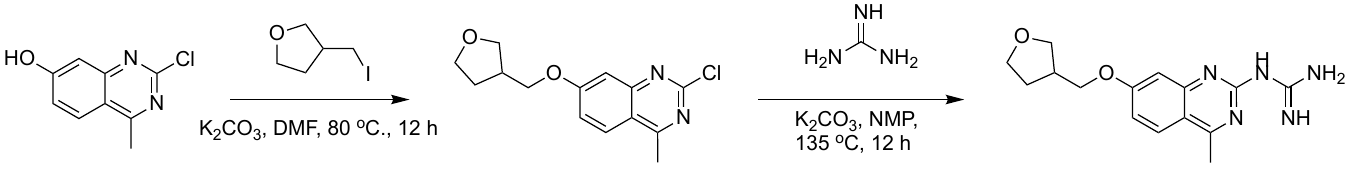

A mixture of 2-chloro-4-methyl-quinazolin-7-ol (0.100 g, 0.514 mmol, 1 eq), 3-(iodomethyl)tetrahydrofuran (163 mg, 0.771 mmol, 1.5 eq), K_2_CO_3_ (213 mg, 1.54 mmol, 3 eq) in DMF (1 mL) was degassed and purged with N_2_ (3X). The mixture was stirred at 80 °C for 12 h under a N_2_ atmosphere. Water was added to the reaction (10 mL), and the mixture was extracted with ethyl acetate (10 mL x 3). The combined organic layers were concentrated under reduced pressure to give a residue. The residue was purified by column chromatography (SiO_2_, petroleum ether/ethyl acetate = 1/0 to 5/1) to furnish 2-chloro-4-methyl-7-(tetrahydrofuran-3-ylmethoxy)quinazoline (70 mg) as a white solid.

A mixture of 2-chloro-4-methyl-7-(tetrahydrofuran-3-ylmethoxy)quinazoline (50 mg, 0.18 mmol, 1 eq), guanidine carbonate (107 mg, 0.592 mmol, 3.3 eq), K_2_CO_3_ (124 mg, 0.897 mmol, 5 eq) in NMP (1 mL) was degassed and purged with N_2_ (3X). The mixture was stirred at 135 °C for 12 h under a N_2_ atmosphere. The mixture was filtered and concentrated under reduced pressure to give a residue. The residue was purified by prep-HPLC (neutral condition, column: Phenomenex C18 80 x 40 mm x 3 mm; mobile phase: [water (NH_4_HCO_3_)-ACN]; B%: 10%-40%,8 min) to give 1-[4-methyl-7-(tetrahydrofuran-3-ylmethoxy)quinazolin-2-yl]guanidine (10 mg) as a white solid. LCMS (M+H)^+^ = 302.1; ^1^H NMR: (400MHz, DMSO-d6) δ 7.86 (d, J = 9.0 Hz, 1H), 7.41-7.07 (m, 2H), 7.00 (d, *J* = 2.2 Hz, 1H), 6.89 (dd, *J* = 2.4, 9.0 Hz, 1H), 4.11-3.96 (m, 2H), 3.84-3.74 (m, 2H), 3.67 (q, *J* = 7.8 Hz, 1H), 3.56 (dd, *J* = 5.6, 8.7 Hz, 1H), 2.77-2.68 (m, 1H), 2.64 (s, 3H), 2.11-2.02 (m, 1H), 1.77-1.67 (m, 1H).

Preparation of TDI-014169

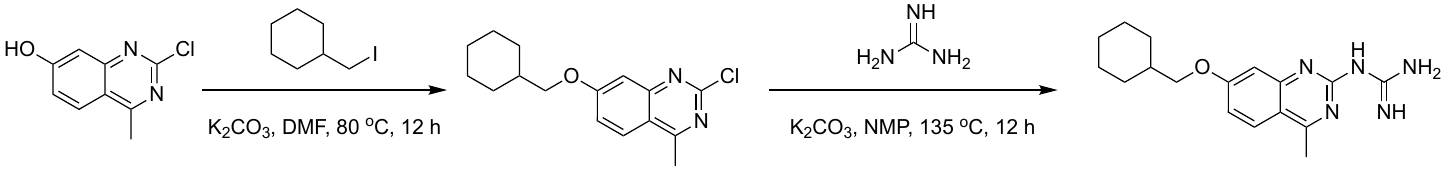

To a solution of 2-chloro-4-methyl-quinazolin-7-ol (100 mg, 0.514 mmol, 1 eq) in DMF (2 mL) was added K_2_CO_3_ (213 mg, 1.54 mmol, 3 eq) and iodomethylcyclohexane (115 mg, 0.514 mmol, 1 eq). The mixture was stirred at 80 °C for 12 h. The mixture was diluted with sat. aqueous NaHCO_3_ (15 mL), and the mixture was extracted with ethyl acetate (10 mL x 3). The combined organic layers were washed with brine (15 mL), dried over Na_2_SO_4_, and filtered. The filtrate was concentrated to furnish 2-chloro-7-(cyclohexylmethoxy)-4-methyl-quinazoline (97 mg, 0.33 mmol, 65 %) as a yellow solid.

To a solution of 2-chloro-7-(cyclohexylmethoxy)-4-methyl-quinazoline (52 mg, 0.18 mmol, 1 eq) in NMP (2 mL) was added K_2_CO_3_ (124 mg, 0.894 mmol, 5 eq) and guanidine carbonate (106 mg, 0.590 mmol, 3.3 eq). The mixture was stirred at 135 °C for 12 h. The reaction was filtered and concentrated. The residue was purified by prep-HPLC (HCl condition) to furnish 1-[7-(cyclohexylmethoxy)-4-methyl-quinazolin-2-yl]guanidine (10 mg, 0.031 mmol, 17%) as a yellow solid. LCMS (M+H)^+^ = 314.1; ^1^H NMR: (400MHz, DMSO-d6) δ 7.71-7.67 (m, 1H), 7.41-7.33 (m, 3H), 7.27-7.20 (m, 4H), 7.17-7.14 (m, 1H), 6.97 (br d, *J* = 8.1 Hz, 1H), 5.13 (s, 1H), 3.97 (br d, *J* = 5.5 Hz, 2H), 3.66-3.62 (m, 3H).

Preparation of TDI-014168

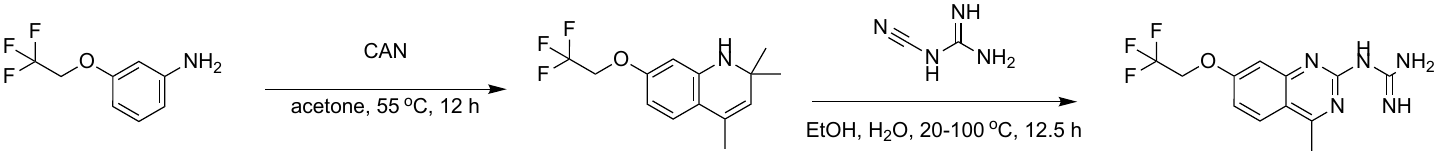

A mixture of 3-(2,2,2-trifluoroethoxy)aniline (330 mg, 1.73 mmol, 1 *eq*), CAN (237 mg, 0.43 mmol, 0.25 *eq*) in acetone (1 mL) was degassed and purged with N_2_ (3X). The mixture was stirred at 55 °C for 12 h under a N_2_ atmosphere. The reaction mixture was concentrated under reduced pressure to remove solvent. The residue was purified by flash silica gel chromatography (ISCO®; 4 g SepaFlash® silica flash column, eluent of 0 to 20% ethyl acetate/petroleum ether gradient at 75 mL/min) to furnish 2,2,4-trimethyl-7-(2,2,2-trifluoroethoxy)-1H- quinoline (150 mg, 0.44 mmol, 26 %) as a white solid.

After stirring a solution of 1-cyanoguanidine (49 mg, 0.58 mmol, 1.2 *eq*) in EtOH (3 mL) and H_2_O (3 mL) at 20 °C for 30 min, 2,2,4-trimethyl-7-(2,2,2-trifluoroethoxy)-1H -quinoline (150 mg, 0.49 mmol, 1 *eq*, HCl) was added. The resulting mixture was stirred at 100 °C for 12 h. The reaction mixture was concentrated under reduced pressure to remove solvent. The mixture was poured into DMSO (4 mL), and the mixture was filtered. The filtrate was concentrated under reduced pressure to furnish 1-[4-methyl-7-(2,2,2-trifluoroethoxy)quinazolin-2-yl]guanidine (72 mg, 0.24 mmol, 50% yield) as a white solid. LCMS (M+H)^+^ = 300.0; ^1^H NMR: (400MHz, MeOH-d4) δ 8.20 (d, *J* = 8.99 Hz, 1H), 7.44 (d, *J* = 2.41 Hz, 1H), 7.33 (dd, *J* = 9.10, 2.30 Hz, 1H), 4.78 (q, *J* = 8.26 Hz, 2H), 2.90 (s, 3H).

Preparation of TDI-012631

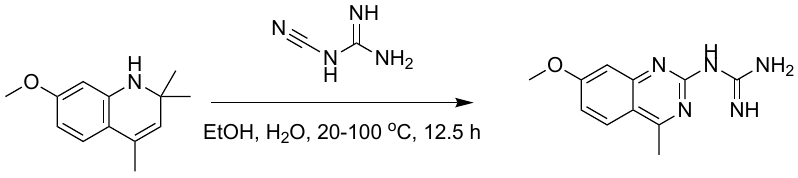

A mixture of 7-methoxy-2,2,4-trimethyl-1H-quinoline (0.100 g, 0.492 mmol, 1 *eq*), 1-cyanoguanidine (62 mg, 0.738 mmol, 1.5 *eq*) in HCl (1 mL 9 N) was degassed and purged with N_2_ (3X). The mixture was stirred at 105 °C for 3 h under a N_2_ atmosphere. The mixture was filtered, and the filter cake was dried under vacuum. The residue was purified by prep-HPLC (HCl condition) to furnish 1-(7-methoxy-4-methyl-quinazolin-2-yl)guanidine (10 mg, 0.046 mmol, 9 %) as a white solid. LCMS (M+H)^+^ = 232.1; ^1^H NMR: (400MHz, DMSO-d6) δ 11.03 (br s, 1H), 8.53 (br s, 3H), 8.14 (d, *J* = 8.99 Hz, 1H), 7.42 (d, *J* = 2.41 Hz, 1H), 7.22 (dd, *J* = 9.10, 2.52 Hz, 1H), 3.94 (s, 3H), 2.83 (s, 3H).

Preparation of TDI-012613

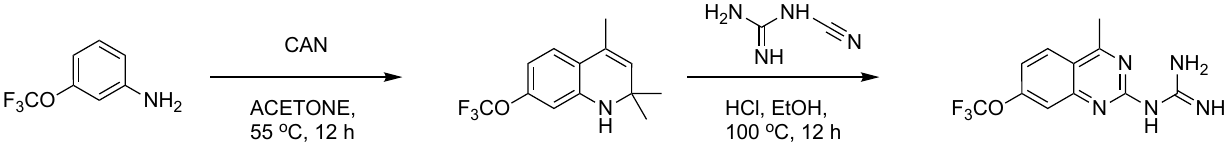

TDI-12613 was prepared in a similar fashion to that described for TDI-12615 using the appropriate aniline in the first step (3-(trifluoromethoxy)aniline). LCMS (M+H)^+^ = 286.0; ^1^H NMR: (400MHz, MeOH-d4) δ 8.39 (d, *J* = 9.2 Hz, 1H), 7.87 (s, 1H), 7.55 (dd, *J* = 2.0, 9.2 Hz, 1 H), 2.99 (s, 3H).

Preparation of TDI-014178

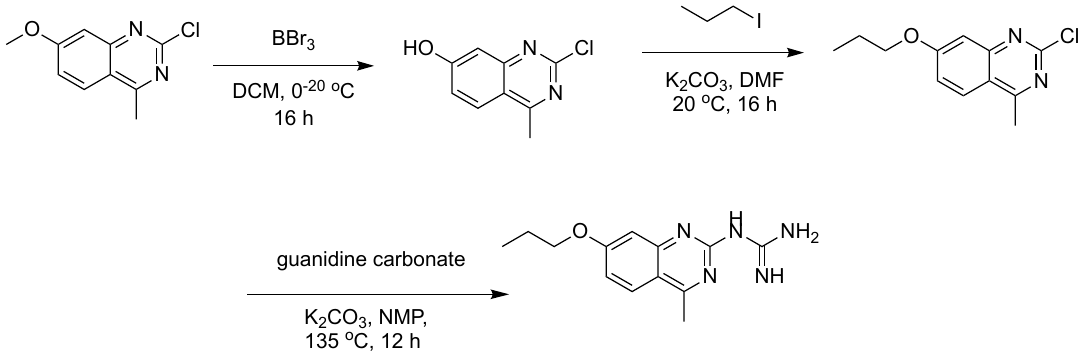

To a solution of 2-chloro-7-methoxy-4-methyl-quinazoline (2.60 g, 12.5 mmol, 1 eq) in DCM (40 mL) was added BBr_3_ (15.6 g, 62.3 mmol, 6.00 mL, 5 eq) at 0 °C. The mixture was stirred at 20 °C for 16 h. The reaction mixture was quenched by addition of ice water (100 mL) at 0 °C. The mixture was filtered and concentrated under reduced pressure to furnish 2-chloro-4-methyl-quinazolin-7-ol (2 g, crude) as a light brown solid.

A mixture of 2-chloro-4-methyl-quinazolin-7-ol (150 mg, 0.771 mmol, 1 eq), 1-iodopropane (200 mg, 1.18 mmol, 1.5 eq), K_2_CO_3_ (213 mg, 1.54 mmol, 2 eq) in DMF (3 mL) was degassed and purged with N_2_ (3X). The mixture was stirred at 20 °C for 16 h under a N_2_ atmosphere. The mixture was poured into H_2_O (50 mL), and the mixture was extracted with ethyl acetate (20 mL x 3). The combined organic layer was washed with brine (50 mL), dried over Na_2_SO_4_, and filtered. The filtrate was concentrated under reduced pressure to give 2-chloro-4-methyl-7-propoxy-quinazoline (0.15 g, crude) as a white solid.

A mixture of 2-chloro-4-methyl-7-propoxy-quinazoline (150 mg, 0.634 mmol, 1 eq), guanidine carbonate (228 mg, 1.27 mmol, 2 eq), K_2_CO_3_ (175 mg, 1.27 mmol, 2 eq) in NMP (1 mL) was degassed and purged with N_2_ (3X). The mixture was stirred at 135 °C for 12 h under a N_2_ atmosphere. The mixture was concentrated under reduced pressure. The residue was purified by prep-HPLC (HCl condition; column: Phenomenex Luna 80 x 30mm x 3 mm; mobile phase: [water(HCl)-ACN];B%: 10%-40%,8 min) to furnish 1-(4-methyl-7-propoxy-quinazolin-2-yl)guanidine (9 mg, 0.03 mmol, 5 %) as a yellow solid. LCMS (M+H)^+^ = 260.1; ^1^H NMR: (400MHz, MeOH-d4) δ 8.10 (d, *J* = 9.1 Hz, 1H), 7.28 (d, *J* = 2.4 Hz, 1H), 7.21 (dd, *J* = 2.4, 9.1 Hz, 1H), 4.21-4.07 (m, 2H), 2.86 (s, 3H), 1.97-1.81 (m, 2H), 1.14-1.04 (m, 3H).

Preparation of TDI-014170

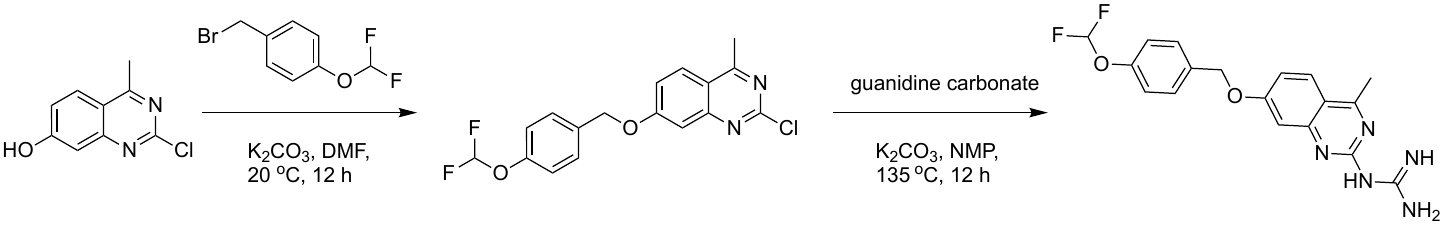

A mixture of 2-chloro-4-methyl-quinazolin-7-ol (0.100 g, 0.514 mmol, 1 eq), 1-(bromomethyl)-4-(difluoromethoxy)benzene (183 mg, 0.771 mmol, 1.5 eq), K_2_CO_3_ (213 mg, 1.54 mmol, 3 eq) in DMF (1.5 mL) was degassed and purged with N_2_ (3X). The mixture was stirred at 20 °C for 12 h under a N_2_ atmosphere. The reaction mixture was partitioned between H_2_O (5 mL) and ethyl acetate (10 mL x 3). The organic phase was washed with brine solution (20 mL), dried over Na_2_SO_4_, and filtered. The filtrate was concentrated under reduced pressure to give a residue. The residue was purified by column chromatography (SiO_2_, petroleum ether/ethyl acetate = 5/1 to 1/1) to furnish 2-chloro-7-[[4-(difluoromethoxy)phenyl]methoxy]-4-methyl-quinazoline (44 mg, 0.13 mmol, 24 %) as a white solid. ^1^H NMR: (400MHz, MeOH-d4) δ 8.23 (dd, *J* = 5.4, 9.2 Hz, 1H), 7.58 (d, *J* = 8.5 Hz, 2H), 7.45-7.40 (m, 2H), 7.27-7.20 (m, 3H), 5.33 (s, 2H), 2.86-2.82 (m, 3H).

A mixture of 2-chloro-7-[[4-(difluoromethoxy)phenyl]methoxy]-4-methyl-quinazoline (0.040 g, 0.11 mmol, 1 eq), guanidine carbonate (68 mg, 0.38 mmol, 3.3 eq), K_2_CO_3_ (79 mg, 0.57 mmol, 5 eq) in NMP (4 mL) was degassed and purged with N_2_ (3X). The mixture was stirred at 135 °C for 12 h under a N_2_ atmosphere. The reaction mixture was concentrated under reduced pressure to remove solvent. The residue was purified by prep-HPLC (neutral condition, column : Phenomenex luna C18 80 x 40 mm x 3 mm; mobile phase: [water (HCl)-ACN]; B%: 28%-45%, 7min) to furnish 1-[7-[[4-(difluoromethoxy)phenyl]methoxy]-4-methyl-quinazolin-2-yl]guanidine (8 mg, 0.02 mmol, 18 %) as a white solid. LCMS (M+H)^+^ = 374.0; ^1^H NMR: (400MHz, MeOH-d4) δ 11.05 (s, 1H), 8.76-8.32 (m, 3H), 8.16 (d, *J* = 9.1 Hz, 1H), 7.59 (s, 1H), 7.57 (s, 1H), 7.52 (d, *J* = 2.4 Hz, 1H), 7.45-7.08 (m, 4H), 5.29 (s, 2H), 2.84 (s, 3H).

Preparation of TDI-012615

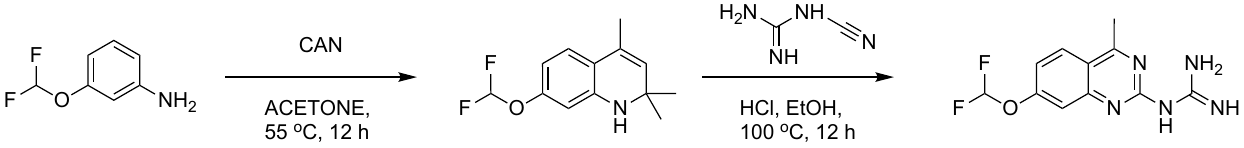

A mixture of 3-(difluoromethoxy)aniline (2.00 g, 12.6 mmol, 1 eq), CAN (1.72 g, 3.14 mmol, 1.57 mL, 0.25 eq) in acetone (20 mL) was degassed and purged with N_2_ (3X). The mixture was stirred at 55 °C for 12 h under a N_2_ atmosphere. The reaction mixture was concentrated under reduced pressure to give 7-(difluoromethoxy)-2,2,4-trimethyl-1H- quinoline (1.00 g, 4.18 mmol, 33 %) as a white solid.

A mixture of 7-(difluoromethoxy)-2,2,4-trimethyl-1H-quinoline (500 mg, 2.09 mmol, 1 eq), 1-cyanoguanidine(176 mg, 2.09 mmol, 1 eq), aq. HCl (2 M, 1.5 mL, 1.44 eq) in EtOH (1.5 mL) was degassed and purged with N_2_ (3X). The mixture was stirred at 100 °C for 1 h under a N_2_ atmosphere. Water (5 mL) was added, and the mixture was extracted with ethyl acetate (15 mL x 3). The combined organic layers were washed with brine (20 mL), dried over Na_2_SO_4_, and filtered. The filtrate was concentrated under reduced pressure to give 1-[7-(difluoromethoxy)-4-methyl- quinazolin-2-yl]guanidine (330 mg, 1.23 mmol, 59 %) as a white solid. LCMS (M+H)^+^ = 268.0; ^1^H NMR: (400MHz, DMSO-d6) δ 11.24-11.12 (m, 1H), 8.67-8.38 (m, 3H), 8.33 (d, J = 9.0 Hz, 1H), 7.76-7.34 (m, 3H), 2.89 (s, 3H).

Preparation of TDI-014173

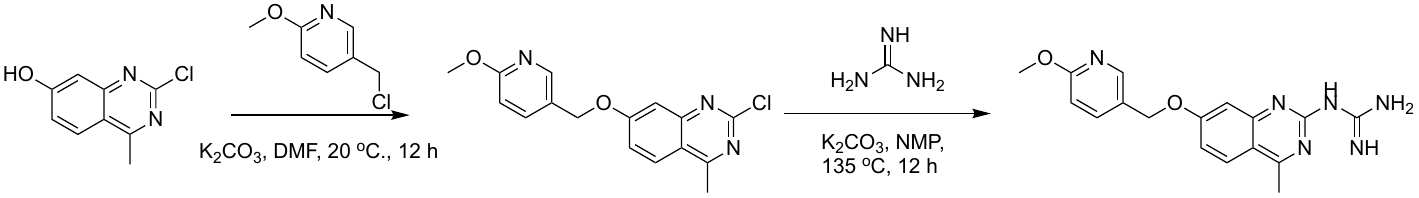

A mixture of 2-chloro-4-methyl-quinazolin-7-ol (100 mg, 0.514 mmol, 1 eq), 5-(chloromethyl)-2-methoxy-pyridine (121 mg, 0.771 mmol, 1.5 eq), K_2_CO_3_ (213 mg, 1.54 mmol, 3 eq) in DMF (1 mL) was degassed and purged with N_2_ (3X). The mixture was stirred at 20 °C for 12 h under a N_2_ atmosphere. Water (10 mL) was added, and the mixture was extracted with ethyl acetate (10 mL x 3). The combined organic layers were dried over Na_2_SO_4_ and filtered. The filtrate was concentrated under reduced pressure to give a residue. The residue was purified by column chromatography (SiO_2_, petroleum ether/ethyl acetate = 1/0 to 5/1) to furnish 2-chloro-7-((6-methoxypyridin-3-yl)methoxy)-4-methylquinazoline (110 mg) as a white solid.

A mixture of 2-chloro-7-((6-methoxypyridin-3-yl)methoxy)-4-methylquinazoline (80 mg, 0.25 mmol, 1 eq), guanidine carbonate (150 mg, 0.836 mmol, 3.3 eq), K_2_CO_3_ (175 mg, 1.27 mmol, 5 eq) in NMP (1 mL) was degassed and purged with N_2_ for (3X). The mixture was stirred at 135 °C for 12 h under a N_2_ atmosphere. The mixture was filtered, and the filtrate was concentrated under reduced pressure to give a residue. The residue was purified by prep-HPLC (neutral condition, column: Phenomenex C18 80 x 40mm x 3 mm; mobile phase: [water (NH_4_HCO_3_)-ACN]; B%: 15%-45%,8min) to furnish 1-(7-((6-methoxypyridin-3-yl)methoxy)-4-methylquinazolin-2-yl)guanidine (18 mg) as a white solid. LCMS (M+H)^+^ = 339.0; ^1^H NMR: (400MHz, DMSO-d6) δ 8.26 (d, *J* = 2.2 Hz, 1H), 8.02 (d, *J* = 9.0 Hz, 1H), 7.83 (dd, *J* = 2.3, 8.7 Hz, 1H), 7.26 (d, *J* = 2.4 Hz, 1H), 7.13 (dd, *J* = 2.4, 8.8 Hz, 1H), 6.86 (d, *J* = 8.6 Hz, 1H), 5.19 (s, 2H), 3.93 (s, 3H), 2.80 (s, 3H).

Preparation of TDI-014171

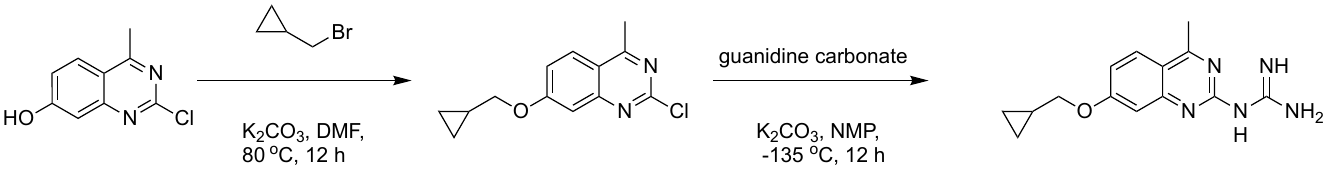

A mixture of 2-chloro-4-methyl-quinazolin-7-ol (108 mg, 0.554 mmol, 1 eq), bromomethylcyclopropane (112 mg, 0.831 mmol, 1.5 eq), K_2_CO_3_ (230 mg, 1.66 mmol, 3 eq) in DMF (1.5 mL) was degassed and purged with N_2_ (3X). The mixture was stirred at 80 °C for 12 h under a N_2_ atmosphere. The reaction mixture was quenched with H_2_O (5 mL), and the mixture was extracted with ethyl acetate (10 mL x 3). The organic phase was separated, washed with brine solution (20 mL), dried over Na_2_SO_4_, and filtered. The filtrate was concentrated under reduced pressure to give a residue. The residue was purified by column chromatography (SiO_2_, petroleum ether/ethyl acetate = 5/1 to 1/1) to furnish 2-chloro-7-(cyclopropylmethoxy)-4-methyl-quinazoline (0.040 g, 0.16 mmol, 29%) as a white solid.

A mixture of 2-chloro-7-(cyclopropylmethoxy)-4-methyl-quinazoline (0.040 g, 0.16 mmol, 1 eq), guanidine carbonate (96 mg, 0.53 mmol, 3.3 eq), K_2_CO_3_ (111 mg, 0.804 mmol, 5 eq) in NMP (4 mL) was degassed and purged with N_2_ (3X). The mixture was stirred at 135 °C for 12 h under a N_2_ atmosphere. The reaction mixture was concentrated under reduced pressure to remove solvent. The residue was purified by prep-HPLC (neutral condition, column: Phenomenex luna C18 80 x 40 mm x 3 mm; mobile phase: [water (HCl)-ACN]; B%: 28%-45%, 7 min) to furnish 1-[7-(cyclopropylmethoxy)-4-methyl-quinazolin-2-yl]guanidine (9 mg, 0.03 mmol, 17 %) as a white solid. LCMS (M+H)^+^ = 272.1; ^1^H NMR: (400MHz, DMSO-d6) δ 11.02 (s, 1H), 8.71-8.30 (m, 3H), 8.13 (d, *J* = 9.1 Hz, 1H), 7.38 (d, *J* = 2.4 Hz, 1H), 7.23 (dd, *J* = 2.5, 9.1 Hz, 1H), 4.01 (d, *J* = 7.0 Hz, 2H), 2.82 (s, 3H), 1.35-1.28 (m, 1H), 0.65-0.60 (m, 2H), 0.41-0.36 (m, 2H).

Preparation of TDI-014175

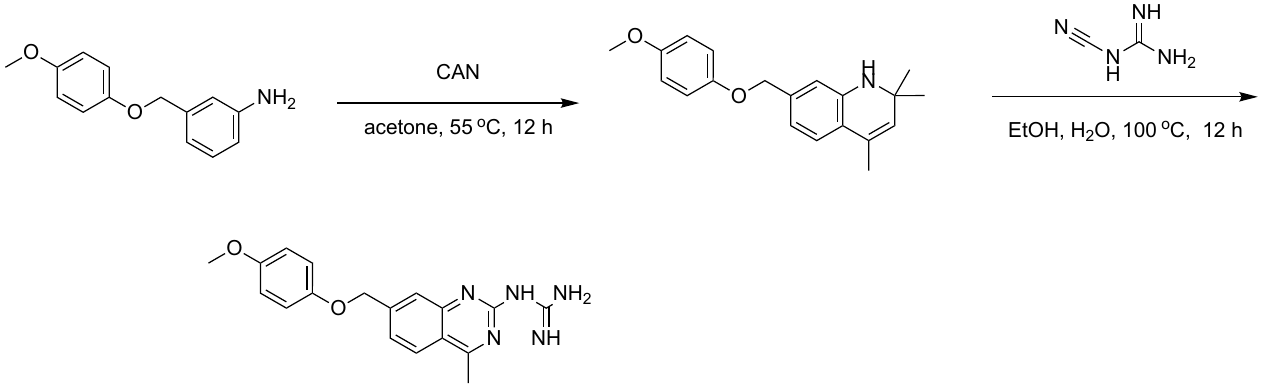

A mixture of 3-[(4-methoxyphenoxy)methyl]aniline (0.600 g, 2.62 mmol, 1 *eq*), CAN (359 mg, 0.654 mmol, 0.25 *eq*) in acetone (5 mL) was degassed and purged with N_2_ (3X). The mixture was stirred at 55 °C for 12 h under a N_2_ atmosphere. The reaction mixture was concentrated under reduced pressure to remove solvent. The residue was purified by flash silica gel chromatography (ISCO®; 12 g SepaFlash® Silica Flash Column, eluent of 0 to 100% ethyl acetate/petroleum ether gradient at 80 mL/min) to furnish 7-[(4-methoxyphenoxy)methyl]-2,2,4-trimethyl-1H-quinoline (384 mg, 1.11 mmol, 42 %, HCl salt) as a gray solid.

To a solution of 7-[(4-methoxyphenoxy)methyl]-2,2,4-trimethyl-1H-quinoline (284 mg, 0.821 mmol, 1 *eq*, HCl) in EtOH (2 mL) and H_2_O (4 mL) was added 1-cyanoguanidine (83 mg, 0.99 mmol, 1.2 *eq*). The mixture was stirred at 100 °C for 12 h. The reaction mixture was concentrated under reduced pressure to remove solvent. The residue was triturated with DMSO at 20 ^o^C for 5 min. The mixture was filtered, and filter cake was dried under reduced pressure to furnish 1-[7-[(4- methoxyphenoxy)methyl]-4-methyl-quinazolin-2-yl]guanidine (22 mg, 0.064 mmol, 8%) as an off-white solid. LCMS (M+H)^+^ = 338.1; ^1^H NMR: (400MHz, DMSO-d6) δ 8.24 (d, J = 8.50 Hz, 1H), 7.98 (s, 1H), 7.70 (dd, J = 8.57, 1.44 Hz, 1H), 7.00-6.94 (m, 2H), 6.88-6.82 (m, 2H), 5.27 (s, 2H), 3.74 (s, 3H), 2.95 (s, 3H).

Preparation of TDI-014184

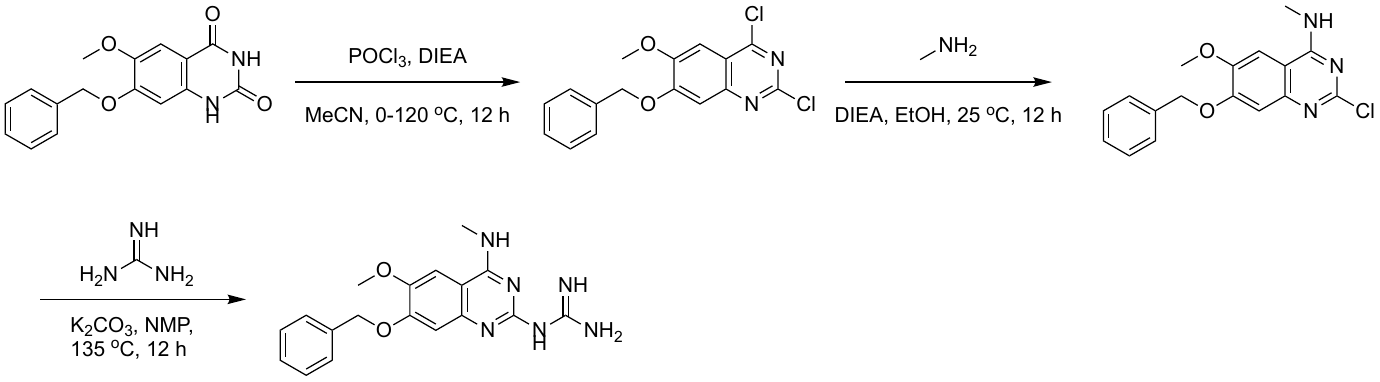

To the mixture of 7-benzyloxy-6-methoxy-1H-quinazoline-2,4-dione (200 mg, 0.670 mmol, 1 eq) in MeCN (5 mL) was added DIPEA (130 mg, 1.01 mmol, 1.5 eq) and POCl_3_ (1.03 g, 6.70 mmol, 10 eq) at 0 °C. The mixture was stirred for 12 h at 120 °C. The mixture was concentrated in vacuo to about 2 mL, and poured in an ice-cold, saturated NaHCO_3_ solution which resulted in the formation of a precipitate. The precipitate that was filtered and dried under vacuum to furnish 7-benzyloxy-2,4-dichloro-6-methoxy-quinazoline (220 mg, crude) as a brown solid.

To the mixture of 7-benzyloxy-2,4-dichloro-6-methoxy-quinazoline (150 mg, 0.448 mmol, 1 eq) in EtOH (2 mL) was added DIPEA (116 mg, 0.895 mmol, 2 eq) and methyl amine-HCl (21 mg, 0.67 umol, 1.5 eq). The mixture was stirred at 25 °C for 12 h. The mixture was poured into H_2_O (10 mL), and the mixture was extracted with ethyl acetate (5 mL x 3). The organic layer was dried with anhydrous Na_2_SO_4_ and filtered. The filtrate was concentrated under vacuum to furnish 7-benzyloxy-2-chloro-6-methoxy-N-methyl-quinazolin-4-amine (127 mg) as a brown solid. LCMS (M+H)^+^ = 330.1.

To the mixture of 7-benzyloxy-2-chloro-6-methoxy-N-methyl-quinazolin-4-amine (70 mg, 0.21 mmol, 1 eq) in NMP (1 mL) was added K_2_CO_3_ (59 mg, 0.42 mmol, 2 eq) and guanidine carbonate (46 mg, 0.25 mmol, 1.2 eq). The mixture was stirred for at 135 °C 12 h. The mixture was filtered and concentrated under vacuum. The residue was purified by pre-HPLC(HCl condition, column: Phenomenex luna C18 80 x 40mm x 3 mm; mobile phase: [water(HCl)-ACN];B%: 18%-38%,7 min) to furnish 1-[7-benzyloxy-6-methoxy-4-(methylamino)quinazolin-2-yl]guanidine (4 mg) as a yellow solid. LCMS (M+H)^+^ = 353.1; ^1^H NMR: (400MHz, DMSO-d6) δ 8.60 (bs, 2H), 8.3 (bs, 3H), 7.48 (s, 1H), 7.44 – 7.37 (m, 5H), 7.28 (s, 1H), 5.21 (s, 2H), 3.86 (s, 3H), 3.00 (bs, 3H).

Preparation of TDI-014182

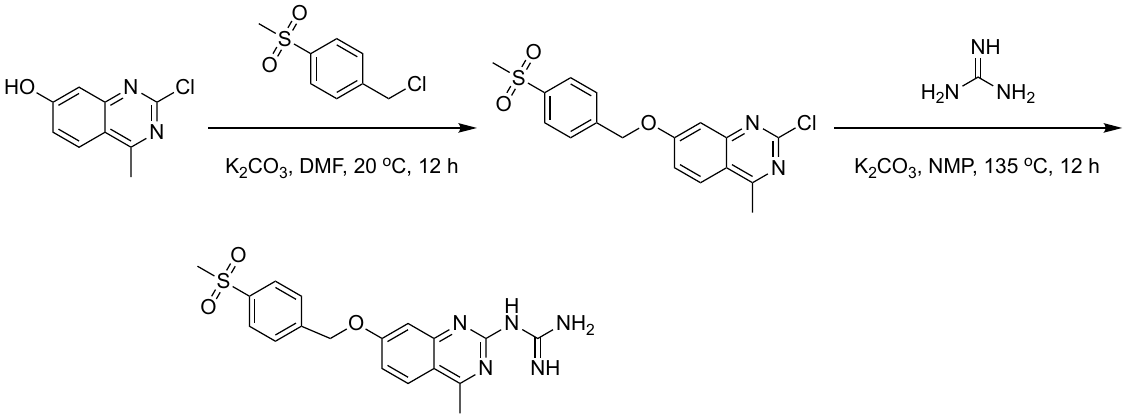

To a solution of 2-chloro-4-methyl-quinazolin-7-ol (200 mg, 1.03 mmol, 1 eq) in DMF (2 mL) was added K_2_CO_3_ (426 mg, 3.08 mmol, 3 eq) and 1-(chloromethyl)-4-methylsulfonyl-benzene (210 mg, 1.03 mmol, 1 eq). The mixture was stirred at 20 °C for 12 h. The reaction was diluted with H_2_O (15 mL), and the mixture was extracted with ethyl acetate (10 mL x 3). The combined organic layers were washed with brine (15 mL), dried over Na_2_SO_4_, and filtered. The filtrate was concentrated under reduced pressure to give a residue. The residue was purified by column chromatography (SiO_2_, petroleum ether/ethyl acetate = 1/0 to 4/1) to furnish 2-chloro-4-methyl-7-[(4-methylsulfonylphenyl)methoxy]quinazoline (236 mg, 0.650 mmol, 63 %) as a yellow solid.

To a solution of 2-chloro-4-methyl-7-[(4-methylsulfonylphenyl)methoxy]quinazoline (100 mg, 0.276 mmol, 1 eq) in NMP (2 mL) was added K_2_CO_3_ (190 mg, 1.38 mmol, 5 eq) and guanidine carbonate (164 mg, 0.910 mmol, 3.3 eq). The mixture was stirred at 135 °C for 12 h. The residue was diluted with H_2_O (10 mL), and the mixture was extracted with EtOAc (10 mL x 3). The combined organic layers were washed with brine (10 mL), dried over Na_2_SO_4_, and filtered. The filtrate was concentrated under reduced pressure to give a residue. The residue was purified by prep-HPLC (neutral condition) to furnish 1-[4-methyl-7-[(4-methylsulfonylphenyl)methoxy]quinazolin-2-yl]guanidine (24 mg, 0.060 mmol, 21 %) was obtained as a yellow solid. LCMS (M+H)^+^ = 386.1; ^1^H NMR: (400MHz, DMSO-d6) δ 11.00 (s, 1H), 8.41 (br s, 3H), 8.20 (d, *J* = 9.1 Hz, 1H), 8.06-7.92 (m, *J* = 8.3 Hz, 2H), 7.85-7.70 (m, *J* = 8.3 Hz, 2H), 7.52 (d, *J* = 2.4 Hz, 1H), 7.34 (dd, *J* = 2.4, 9.2 Hz, 1H), 5.46 (s, 2H), 3.28-3.16 (m, 3H), 2.85 (s, 3H).

Preparation of TDI-014188

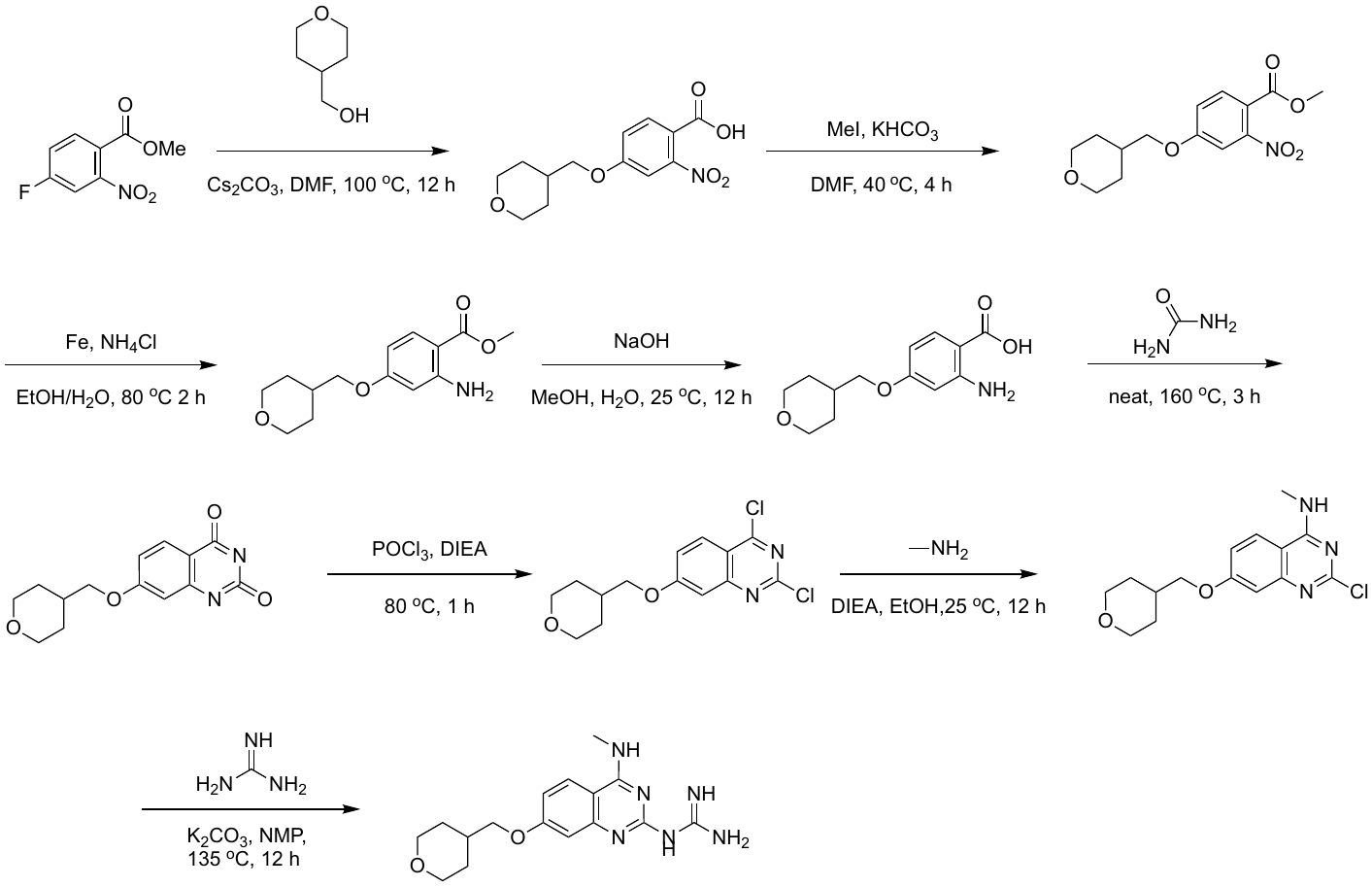

To the mixture of methyl 4-fluoro-2-nitro-benzoate (3.00 g, 15.1 mmol, 1 eq) in DMF (30 mL) was added tetrahydropyran-4-ylmethanol (2.10 g, 18.1 mmol, 1.2 eq) and Cs_2_CO_3_ (14.7 g, 45.2 mmol, 3 eq). The mixture was stirred at 100 °C for 12 h. The mixture was filtered and concentrated in vacuum. Then the residue was diluted with H_2_O (10 mL). The pH of the mixture was adjusted to 3 by addition of aqueous HCl (2 N). The yellow precipitate was collected and dried to furnish 2-nitro-4-(tetrahydropyran-4-ylmethoxy)benzoic acid (3.53 g, crude) as a yellow solid.

To the mixture of 2-nitro-4-(tetrahydropyran-4-ylmethoxy)benzoic acid (2.50 g, 8.89 mmol, 1 eq) in DMF (25 mL) was added KHCO_3_ (1.07 g, 10.7 mmol, 1.2 eq) and MeI (1.89 g, 13.3 mmol, 1.5 eq). The mixture was stirred at 40 °C for 4 h. The mixture was quenched with H_2_O (10 mL), and the mixture was exacted with ethyl acetate (10 mL x 3). The combined organic layer was washed with brined, dried with anhydrous Na_2_SO_4_, and filtered. The filtrate was concentrated under vacuum. The residue was purified by silica gel chromatography (petroleum ether/ethyl acetate = 5/1 to 1/1) to furnish methyl 2-nitro-4-(tetrahydropyran-4-ylmethoxy) benzoate (1.58 g, crude) as a yellow solid.

To the mixture of methyl 2-nitro-4-(tetrahydropyran-4-ylmethoxy)benzoate (1.58 g, 5.35 mmol, 1 eq) in EtOH (15 L) and H_2_O (15 mL) was added Fe (1.49 g, 26.8 mmol, 5 eq) and NH_4_Cl (2.86 g, 53.5 mmol, 10 eq). The mixture was stirred at 80 °C for 2 h. The mixture was extracted with ethyl acetate (20 mL x 3). The combined organic layers were dried with anhydrous Na_2_SO_4_ and filtered. The filtrate was concentrated under vacuum to furnish methyl 2-amino-4-(tetrahydropyran-4-ylmethoxy)benzoate (749 mg, 2.82 mmol, 52%) as a yellow solid.

To the mixture of methyl 2-amino-4-(tetrahydropyran-4-ylmethoxy)benzoate (70 mg, 1.51 mmol, 1 eq) in MeOH (4 mL) and H_2_O (5 mL) was added NaOH (166 mg). The mixture was stirred for 12 h at 25 °C. The mixture was poured into H_2_O (5 ml), and the pH of the mixture was adjusted to 3 by addition of aqueous HCl (2 N). The brown precipitate was collected and dried to furnish 2-amino-4-(tetrahydropyran-4-ylmethoxy)benzoic acid (220 mg, 0.876 mmol, 58%) as a yellow solid.

A mixture of 2-amino-4-(tetrahydropyran-4-ylmethoxy)benzoic acid (420 mg, 1.67 mmol, 1 eq), urea (602 mg, 10.0 mmol, 6 eq) was degassed and purged with N_2_ (3X). The mixture was stirred at 160°C for 3 h under a N_2_ atmosphere. The mixture was triturated with H_2_O (10 mL) and acetone (10 mL). The formed solid was collected and dried to furnish 7-(tetrahydropyran-4-ylmethoxy)-1H-quinazoline-2,4-dione (80 mg, 0.29 mmol, 17 %) as a brown solid.

A mixture of 7-(tetrahydropyran-4-ylmethoxy)-1H-quinazoline-2,4-dione (80 mg, 0.29 mmol, 1 eq), POCl_3_ (444 mg, 2.90 mmol, 10 eq), DIPEA (150 mg, 1.16 mmol, 4 eq) was degassed and purged with N_2_ (3X). The mixture was stirred at 80 °C for 1 h under a N_2_ atmosphere. The mixture was concentrated under vacuum. The residue was purified by prep-TLC (SiO_2_, petroleum ether/ethyl acetate = 3/1) to furnish 2,4-dichloro-7-(tetrahydropyran-4-ylmethoxy)quinazoline (40 mg, 0.12 mmol, 42%) as a white solid.

To the mixture of 2,4-dichloro-7-(tetrahydropyran-4-ylmethoxy)quinazoline (34 mg, 0.11 mmol, 1 eq) in EtOH (1 mL) was added DIPEA (28 mg, 0.22 mmol, 2 eq) and methylamine-HCl (9 mg, 0.13 mmol, 1.2 eq). The mixture was stirred at 25 °C for 12 h. The mixture was extracted ethyl acetate (2 mL x 3). The combined organic layers were dried with anhydrous Na_2_SO_4_ and filtered. The filtrate was concentrated under vacuum to furnish 2-chloro-N-methyl-7-(tetrahydropyran-4-ylmethoxy)quinazolin-4-amine (40 mg) as a white solid.

To the mixture of 2-chloro-N-methyl-7-(tetrahydropyran-4-ylmethoxy)quinazolin-4-amine (34 mg, 0.11 mmol, 1 eq) in NMP (1 mL) was added K_2_CO_3_ (46 mg, 0.33 mmol, 3 eq) and guanidine carbonate (22 mg, 0.12 mmol, 1.1 eq). The mixture was stirred at 135 °C for 12 h. The mixture was filtered and concentrated in vacuum. The residue was purified by Prep-HPLC (HCl condition, column: Phenomenex Luna 80 x 30mm x 3 mm; mobile phase:[water(HCl)-ACN]; B%: 1%-30%, 8 min) to furnish 1-[4-(methylamino)-7-(tetrahydropyran-4-ylmethoxy)quinazolin-2-yl]guanidine (11 mg, 0.033 mmol, 30 %) as a white solid. LCMS (M+H)^+^ = 331.1; ^1^H NMR: (400MHz, DMSO-d6) δ 8.70 (bs, 1H), 8.12 (d, J = 9.2 Hz, 1H), 7.13 (d, J = 2.4 Hz, 1H), 7.02 (m, 1H), 3.90 (m, 2H), 3.88 (m, 2H), 3.35 (m, 2H), 2.98 (d, J = 4.4 Hz, 3H), 2.05 (m, 1H), 1.68-1.71 (m, 2H), 1.39-1.33 (m, 2H).

Preparation of TDI-012943

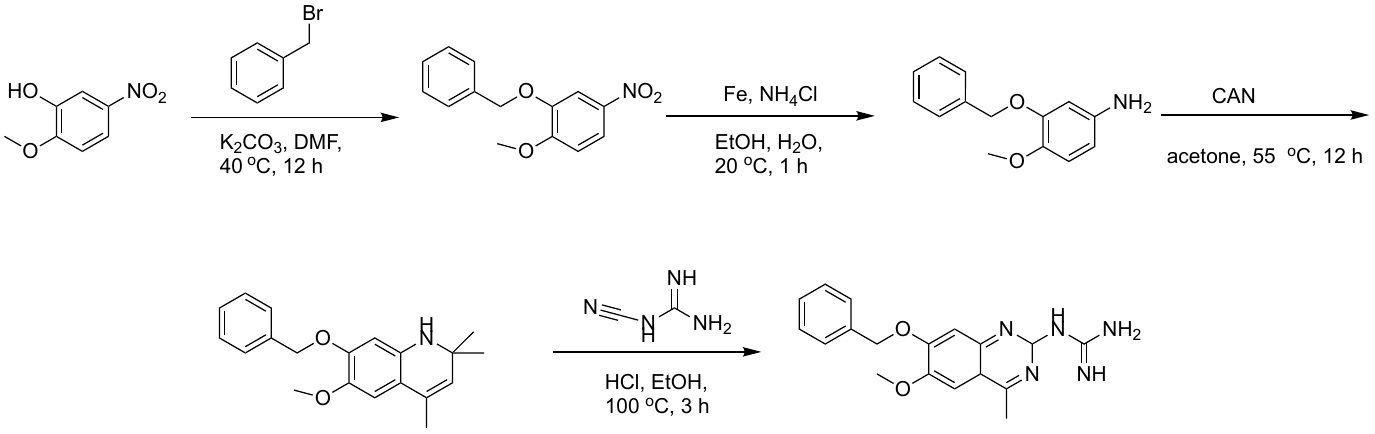

To the mixture of 2-methoxy-5-nitro-phenol (5.00 g, 29.6 mmol, 1 *eq*) in DMF (6 mL) was added benzyl bromide (5.56 g, 32.5 mmol, 3.86 mL, 1.1 *eq*) and K_2_CO_3_ (6.13 g, 44.3 mmol, 1.5 *eq*) under N_2_. The mixture was stirred at 40°C for 12 h. The mixture was diluted with H_2_O (20 mL), and the mixture was extracted with ethyl acetate (10 mL x 3). The combined organic phase was dried with anhydrous Na_2_SO_4_ and filtered. The filtrate was concentrated in vacuum. The residue was purified by silica gel chromatography (petroleum ether/ethyl acetate = 50/1 to 1/1) to furnish 2-benzyloxy-1-methoxy-4-nitro-benzene (5.90 g, 22.8 mmol, 77 %) as white solid.

A mixture of 2-benzyloxy-1-methoxy-4-nitro-benzene (1.00 g, 3.86 mmol, 1 *eq*), Fe (1.08 g, 19.3 mmol, 5 *eq*) and NH_4_Cl (1.03 g, 19.3 mmol, 5 *eq*) in EtOH (5 mL) and H_2_O (5 mL) was stirred at 20 °C for 1 h. The mixture was filtered, and the filtrate was extracted with ethyl acetate (5 mL x 3). The combined organic phase was washed with brine, dried with anhydrous Na_2_SO_4_, and filtered. The filtrate was concentrated under vacuum. The residue was purified by silica gel chromatography (petroleum ether/ethyl acetate = 50/1 to 1/1) to furnish 3-benzyloxy-4-methoxy-aniline (0.800 g, 3.49 mmol, 90 %) as yellow solid.

A mixture of 3-benzyloxy-4-methoxy-aniline (0.200 g, 0.872 mmol, 1 *eq*) and CAN (120 mg, 0.218 mmol, , 0.25 *eq*) in acetone (3 mL) was stirred at 55°C for 12 h. The mixture was concentrated under vacuum to furnish 7-benzyloxy-6-methoxy-2,2,4-trimethyl-1H-quinoline (0.170 g, 0.550 mmol, 63%) as yellow solid.

To a solution of 7-benzyloxy-6-methoxy-2,2,4-trimethyl-1H-quinoline (120 mg, 0.388 mmol, 1 eq) in EtOH (2 mL) was added aqueous HCl (1 M, 0.39 mL, 1 eq) and 1-cyanoguanidine (33 mg, 0.39 mmol, 1 eq). The mixture was stirred at 100 °C for 3 h. The mixture was concentrated under vacuum. The residue was purified by prep-HPLC (column: Welch Xtimate C18 100 x 25mm x 3 mm; mobile phase: [water(0.04%HCl)-ACN];B%: 10%-30%,8 min) to furnish 1-(7-benzyloxy-6-methoxy-4-methyl-quinazolin-2-yl)guanidine (52 mg) as white solid. LCMS (M+H)^+^ = 338.1; ^1^H NMR: (400MHz, DMSO-d6) 10.92 (s, 1H), 8.28 (bs, 3H), 7.55 – 7.42 (m, 7H), 5.30 (s, 2H), 3.96 (s, 3H), 2.85 (s, 3H).

Preparation of TDI-013000

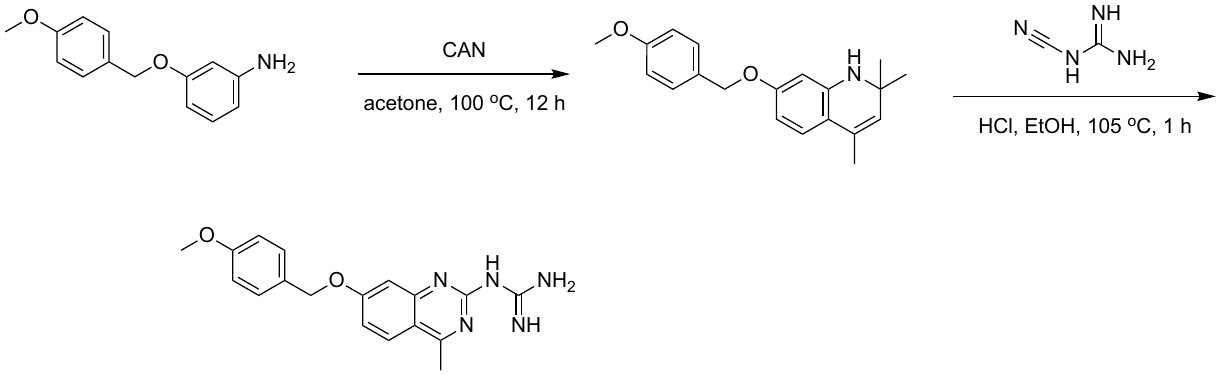

To a solution of 3-[(4-methoxyphenyl)methoxy]aniline (1.40 g, 6.11 mmol, 1 eq) in acetone (25 mL) was added CAN (837 mg, 1.53 mmol, 0.25 eq). The mixture was stirred at 100 °C for 12 h. The mixture was concentrated under reduced pressure to give a residue. The residue was purified by column chromatography (SiO_2_, petroleum ether/ethyl acetate = 1/0 to 9/1) to furnish 7-[(4-methoxyphenyl)methoxy]-2,2,4-trimethyl-1H-quinoline (0.400 g, 1.29 mmol, 21%) as a yellow solid.

To a solution of 7-[(4-methoxyphenyl)methoxy]-2,2,4-trimethyl-1H-quinoline (0.050 g, 0.16 mmol, 1 eq) in EtOH (0.75 mL) was added aqueous HCl (1 M, 0.16 mL, 1 eq) and 1-cyanoguanidine (14 mg, 0.16 mmol, 1 eq). The mixture was stirred at 105 °C for 1 h. The residue was purified by prep-HPLC (HCl condition) to furnish 1-[7-[(4-methoxyphenyl)methoxy]-4-methyl-quinazolin-2-yl]guanidine (0.030 g, 0.089 mmol, 55%) as a white solid. LCMS (M+H)^+^ = 338.1; ^1^H NMR: (400MHz, DMSO-d6) δ 11.01 (s, 1H), 8.46 (br s, 2H), 8.16 (d, *J* = 9.2 Hz, 1H), 7.52 (d, *J* = 2.4 Hz, 1H), 7.45 (d, *J* = 8.8 Hz, 2H), 7.27 (dd, *J* = 2.5, 9.1 Hz, 1H), 7.00-6.96 (m, 2H), 5.21 (s, 2H), 3.78-3.76 (m, 3H), 2.84 (s, 3H).

Preparation of TDI-014190

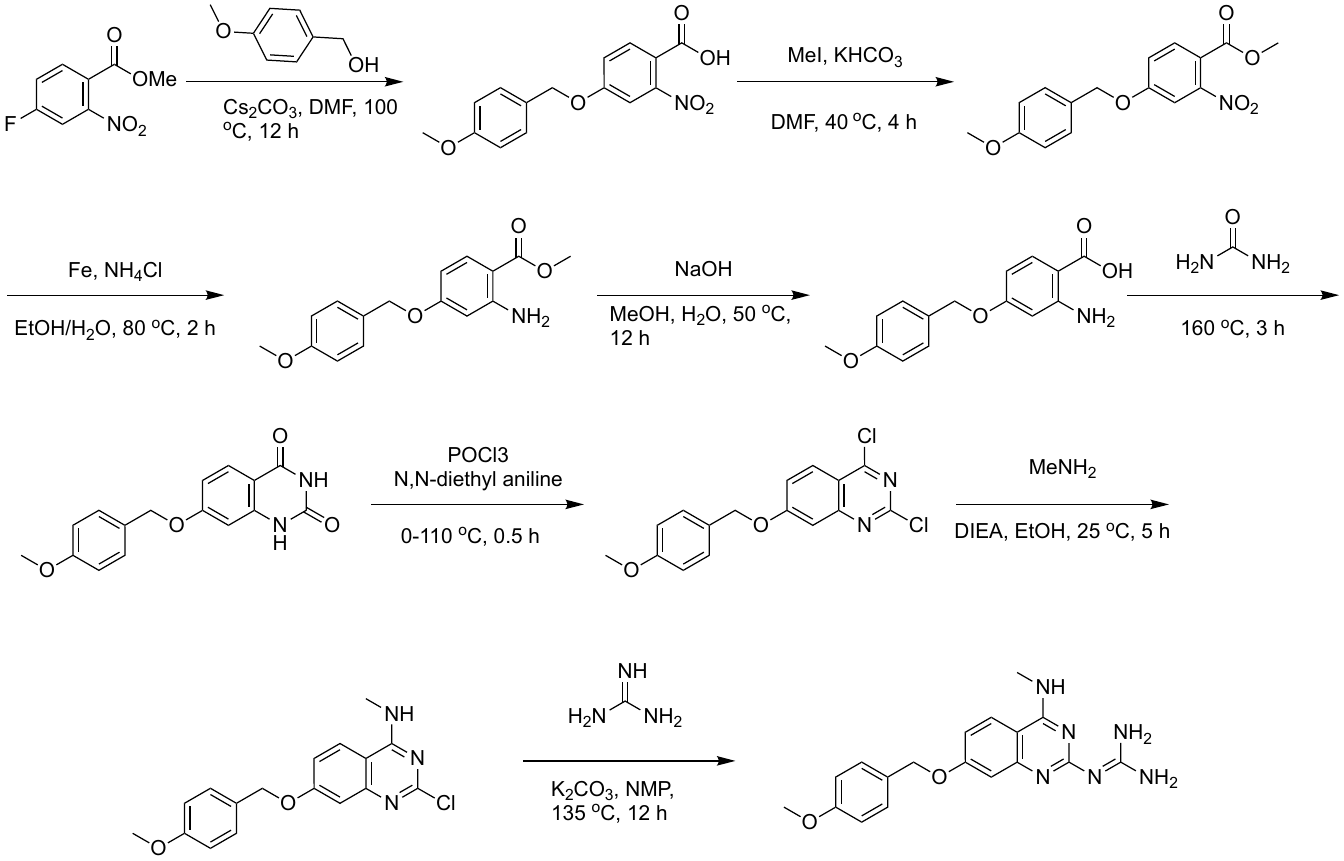

To the mixture of methyl 4-fluoro-2-nitro-benzoate (6.00 g, 30.1 mmol, 1 eq) in DMF (60 mL) was added (4-methoxyphenyl)methanol (5.00 g, 36.2 mmol, 4.50 mL, 1.2 eq) and Cs_2_CO_3_ (29.5 g, 90.4 mmol, 3 eq). The mixture was stirred at 100 °C for 12 h. The mixture filtered and concentrated under vacuum. The mixture was extracted with ethyl acetate (30 mL x 3), and the pH of the water phase was adjusted to 3 by addition of aqueous HCl (2 N). The yellow precipitate was collected in a Büchner funnel under suction filtration and dried to furnish 4-((4-methoxybenzyl)oxy)-2-nitrobenzoic acid (2.38 g, 7.73 mmol, 26%) as a yellow solid.

To the mixture of 4-((4-methoxybenzyl)oxy)-2-nitrobenzoic acid (2.35 g, 7.75 mmol, 1 eq) in DMF (25 mL) was added KHCO_3_ (931 mg, 9.30 mmol, 1.2 eq) and MeI (1.65 g, 11.6 mmol, 1.5 eq). The mixture was stirred at 40 °C for 4 h. The mixture was extracted with ethyl acetate (20 mL x 3). The organic layer was dried with anhydrous Na_2_SO_4_, and filtered. The filtrate was concentrated under vacuum. The residue was purified by silica gel chromatography(petroleum ether/ethyl acetate = 5/1 to 1/1) to furnish methyl 4-((4-methoxybenzyl)oxy)-2-nitrobenzoate (2.36 g, 6.90 mmol, 89 %) as a yellow solid.

To the mixture of methyl 4-((4-methoxybenzyl)oxy)-2-nitrobenzoate (1.10 g, 3.47 mmol, 1 eq) in H_2_O (15 mL) and EtOH (15 mL) was added Fe (968 mg, 17.3 mmol, 5 eq) and NH_4_Cl (1.85 g, 34.7 mmol, 10 eq). The mixture was stirred at 80 °C for 2 h. The mixture was extracted with ethyl acetate (30 mL x 3). The organic layer was dried with anhydrous Na_2_SO_4_ and filtered. The filtrate was concentrated under vacuum to furnish methyl 2-amino-4-((4-methoxybenzyl)oxy) benzoate (1 g, crude) as a yellow solid.

To the mixture of methyl 2-amino-4-((4-methoxybenzyl)oxy)benzoate (1.00 g, 3.48 mmol, 1 eq) in MeOH (12 mL) and H_2_O (12 mL) was added NaOH (278 mg, 6.96 mmol, 2 eq). The mixture was stirred at 50 °C for 12 h. The mixture was extracted with ethyl acetate (20 mL x 3), and the pH of the water phase was adjusted to 3 by addition of aqueous HCl (2 N). The water phase was extracted with ethyl acetate (10 mL x 3). The organic layer was dried with anhydrous Na_2_SO_4_ and filtered. The filtrate was concentrated under vacuum to furnish 2-amino-4-((4-methoxybenzyl)oxy)benzoic acid (643 mg, crude) as a brown solid.

A mixture of 2-amino-4-((4-methoxybenzyl)oxy)benzoic acid (640 mg, 2.34 mmol, 1 eq), urea (844 mg, 14.1 mmol, 6 eq) was degassed and purged with N_2_ (3X). The mixture was stirred at 160 ^o^C for 3 h under a N_2_ atmosphere. The mixture was triturated with H_2_O (10 mL) and acetone (10 mL). The formed solid was collected and dried to furnish 7-((4-methoxybenzyl)oxy) quinazoline-2,4(1H,3H)-dione (193 mg, 0.647 mmol, 28%) a white solid.

To a mixture of 7-((4-methoxybenzyl)oxy)quinazoline-2,4(1H,3H)-dione (60 mg, 0.20 mmol, 1 eq) in *N*,*N*-diethylaniline (0.2 mL) at 0 °C was added dropwise POCl_3_ (0.4 mL). The mixture was stirred at 110 °C for 0.5 h. The mixture was filtered and concentrated in vacuum. The residue was purified by silica gel chromatography (petroleum ether/ethyl acetate = 10/1 to 1/1) to furnish 2,4-dichloro-7-((4-methoxybenzyl)oxy)quinazoline (96 mg) as a yellow solid.

To the mixture of 2,4-dichloro-7-((4-methoxybenzyl)oxy)quinazoline (80 mg, 0.24 mmol, 1 eq) in EtOH (1 mL) was added DIPEA (62 mg, 0.48 mmol, 2 eq) and methyl amine-HCl (19 mg, 0.29 mmol, 1.2 eq). The mixture was stirred at 25 °C for 5 h. The mixture was extracted with ethyl acetate (2 mL x 3). The organic layer was dried with anhydrous Na_2_SO_4_ and filtered. The filtrate was concentrated under vacuum to furnish 2-chloro-7-((4-methoxybenzyl)oxy)-N-methylquinazolin-4-amine (91 mg, crude) as a yellow solid.

To the mixture of 2-chloro-7-((4-methoxybenzyl)oxy)-N-methylquinazolin-4-amine (70 mg, 0.21 mmol, 1 eq) in NMP (1 mL) was added K_2_CO_3_ (88 mg, 0.64 mmol, 3 eq) and guanidine carbonate (46 mg, 0.25 mmol, 1.2 eq). The mixture was stirred at 135 °C for 12 h. The mixture was filtered and concentrated under vacuum. The mixture was purified by prep-HPLC (column: Phenomenex Luna 80 x 30mm x 3 mm; mobile phase:[water(HCl)-ACN];B%: 1%-25%, 8 min) to furnish 2-(7-((4-methoxybenzyl)oxy) -4-(methylamino)quinazolin-2-yl)guanidine (29 mg, 0.079 mmol, 37%) as a white solid. LCMS (M+H)^+^ = 353.1; ^1^H NMR: (400MHz, DMSO-d6) δ 8.69-8.64 (m, 1H), 8.64-8.642 (m, 1H), 8.27 (d, *J* = 3.2 Hz, 1H), 7.41 (d, *J* = 8.4 Hz, 2H), 7.22-7.20 (m, 1H), 7.07 (d, *J* = 9.2 Hz, 1H), 7.07 (d, *J* = 40.4 Hz, 2H), 5.13 (s, 2H), 3.76 ( s, 3H), 2.98 (d, *J* = 4.4 Hz, 3H).

Preparation of TDI-014185

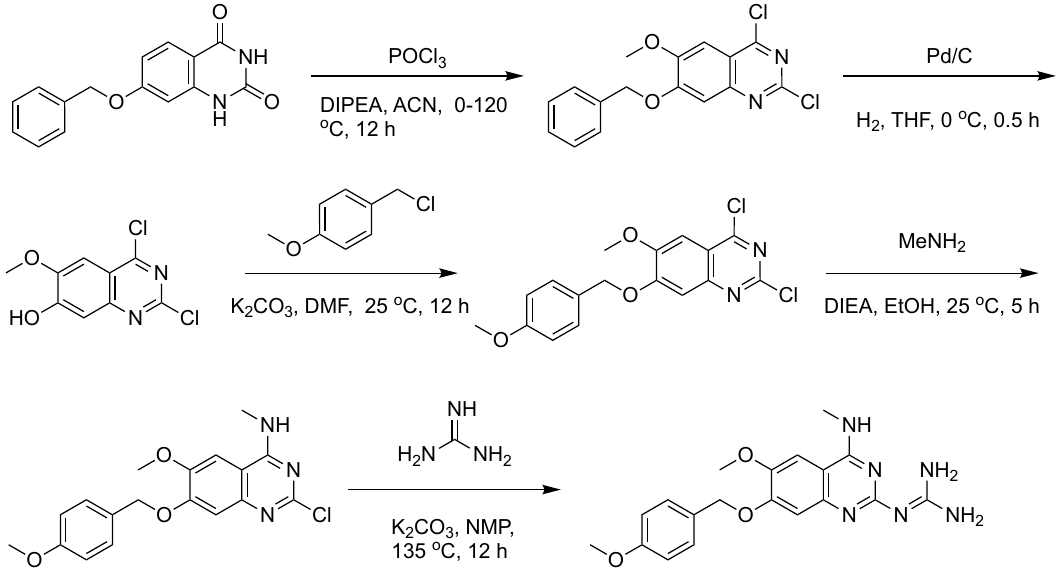

To the mixture of 7-benzyloxy-6-methoxy-1H-quinazoline-2,4-dione (1.80 g, 6.0 mmol, 1 eq) in MeCN (40 mL) was added DIPEA (1.17 g, 9.05 mmol, 1.58 mL, 1.5 eq) and POCl_3_ (18.5 g, 121 mmol, 11.22 mL, 20 eq) at 0 °C. The mixture was stirred at 120 °C for 12 h. The mixture was concentrated under vacuo to about 5 mL. The residue was poured in an ice-cold saturated NaHCO_3_ solution. The formed precipitate was collected and dried to furnish 7-(benzyloxy)-2,4-dichloro-6-methoxyquinazoline (1.62 g, crude) as a brown solid.

To a solution of 7-(benzyloxy)-2,4-dichloro-6-methoxyquinazoline (300 mg, 0.895 mmol, 1 eq) in THF (6 mL) was added Pd/C (50 mg, 10 wt% on carbon) under H_2_ . The suspension was degassed under vacuum and purged with H_2_ several times. The mixture was stirred under H_2_ (15 psi) at 0 °C for 0.5 h. The mixture was filtered and concentrated under vacuum to furnish 2,4-dichloro-6-methoxyquinazolin-7-ol (650 mg, crude) as a yellow solid.

To the mixture of 2,4-dichloro-6-methoxyquinazolin-7-ol (150 mg, 0.612 mmol, 1 eq) in DMF (2 mL) was added K_2_CO_3_ (169 mg, 1.22 mmol, 2 eq) and 1-(chloromethyl)-4-methoxy-benzene (115 mg, 0.735 mmol, 1.2 eq). The mixture was stirred at 25 °C for 2 h. The mixture was extracted with ethyl acetate (5 mL x 3). The organic layer was dried with anhydrous Na_2_SO_4_ and filtered. The filtrate was concentrated under vacuum. The residue was purified by silica gel chromatography (petroleum ether/ethyl acetate = 3/1 to 1/1) to furnish 2,4-dichloro-6-methoxy-7-((4-methoxybenzyl)oxy) quinazoline (50 mg, crude) as a yellow solid.

To the mixture of 2,4-dichloro-6-methoxy-7-((4-methoxybenzyl)oxy)quinazoline (80 mg, 0.22 mmol, 1 eq) in EtOH (1 mL) was added DIPEA (57 mg, 0.44 mmol, 2 eq) and methylamine-HCl (22 mg, 0.33 mmol). The mixture was stirred at 25 °C for 12 h. The mixture was extracted with ethyl acetate (5 mL x 3). The organic layer was dried with anhydrous Na_2_SO_4_ and filtered. The filtrate was concentrated under vacuum to furnish 2-chloro-6-methoxy-7-((4-methoxybenzyl)oxy)-N-methylquinazolin-4-amine (66 mg, crude) as a yellow solid.

To a solution of 2-chloro-6-methoxy-7-((4-methoxybenzyl)oxy)-N-methylquinazolin-4- amine (60 mg, 0.17 mmol, 1 eq) in NMP (1 mL) was added K_2_CO_3_ (46 mg, 0.33 mmol, 2 eq) and guanidine carbonate (45 mg, 0.25 mmol, 1.5 eq). The mixture was stirred at 135 °C for 12 h. The mixture was filtered and concentrated under vacuum. The residue was purified by prep-HPLC(HCl condition, column: Phenomenex Luna 80 x 30mm x 3 mm; mobile phase: [water(HCl)-ACN];B%: 5%-30%, 8 min) to furnish 2-(6-methoxy-7-((4-methoxybenzyl)oxy)-4-(methylamino)quinazolin-2-yl) guanidine (5 mg, HCl salt) as a white solid. LCMS (M+H)^+^ = 383.1; ^1^H NMR: (400MHz, DMSO-d6) δ 8.76-7.97 (m, 5H), 7.67 (s, 1H), 7.42 (d, *J* = 8.6 Hz, 2H), 7.28 (s, 1H), 6.97 (d, *J* = 8.5 Hz, 2H), 5.11 (s, 2H), 3.85 (s, 3H), 3.77 (s, 3H), 3.00 (br d, *J* = 4.0 Hz, 3H).

Preparation of TDI-014180

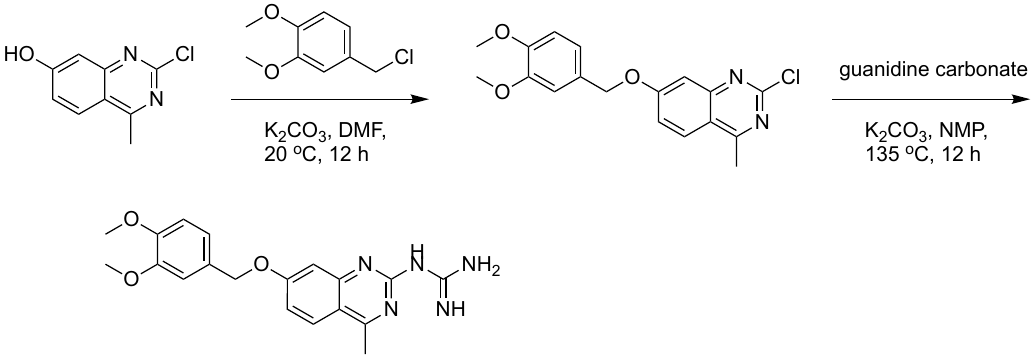

To a solution of 2-chloro-4-methyl-quinazolin-7-ol (130 mg, 0.668 mmol, 1 eq) in DMF (2.5 mL) was added K_2_CO_3_ (277 mg, 2.00 mmol, 3 eq) and 4-(chloromethyl)-1,2-dimethoxy-benzene (130 mg, 0.697 mmol, 1.04 eq) . The mixture was stirred at 25 °C for 12 h. Water (20 mL) was added, and the mixture was extracted with ethyl acetate (10 mL x 3). The combined organic layers were washed with brine (20 mL), dried over Na_2_SO_4_, and filtered. The filtrate was concentrated under reduced pressure to give a residue. The residue was purified by column chromatography (SiO_2_, petroleum ether/ethyl acetate = 9/1 to 1/1) to furnish 2-chloro-7-[(3,4-dimethoxyphenyl)methoxy]-4-methyl-quinazoline (130 mg, crude) as a white solid.

To a solution of 2-chloro-7-[(3,4-dimethoxyphenyl)methoxy]-4-methyl-quinazoline (100 mg, 0.290 mmol, 1 eq) in NMP (2 mL) was added K_2_CO_3_ (120 mg, 0.870 mmol, 3 eq) and guanidine carbonate (57 mg, 0.32 mmol, 1.1 eq) . The mixture was stirred at 135 ^o^C for 12 h. The reaction mixture was concentrated under reduced pressure. The residue was purified by prep-HPLC (formic acid (FA) conditions, column: Phenomenex Luna C18 150 x 30mm x 5 mm; mobile phase: [water(FA)-ACN];B%: 20%-50%,8min) to furnish 1-[7-[(3,4-dimethoxyphenyl)methoxy]-4-methyl-quinazolin-2-yl]guanidine (4 mg, 0.01 mmol, 3%, formic acid salt) as a white solid. LCMS (M+H)^+^ = 368.1; ^1^H NMR: (400MHz, DMSO-d6) δ 8.86-8.49 (m, 3H), 8.43 (s, 1H), 8.06 (d, *J* = 9.1 Hz, 1H), 7.41 (br d, *J* = 1.9 Hz, 1H), 7.17 (dd, *J* = 2.4, 9.0 Hz, 1H), 7.12 (d, J=1.5 Hz, 1H), 7.07-7.01 (m, 1H), 7.00-6.94 (m, 1H), 5.18 (s, 2H), 3.77 (d, *J* = 4.9 Hz, 6H), 2.77 (s, 3H).
